# AI-enabled Spatial Profiling of Circulating Tumor-Immune Ecosystems Predicts Patient Outcomes Across Cancers

**DOI:** 10.64898/2026.07.02.736133

**Authors:** Joshua R. Squires, Yuanfei Sun, Andrew Hoffmann, Youbin Zhang, Hao Pan, Fangjia Tong, Yiwen He, David Scholten, Hannah Almubarak, Michael Gurley, Allegra Minor, Anmol Singh, James Zhang, Hannah Ding, Chengsheng Mao, Leonidas C. Platanias, Jindan Yu, Maha Hussain, Yuan Luo, William J. Gradishar, Massimo Cristofanilli, Lee A.D. Cooper, Lili Zhao, Deyu Fang, Carsen Stringer, Huiping Liu

**Author notes:** Co-first authors with equal contributions (Joshua R. Squires, Yuanfei Sun, Andrew Hoffmann, Youbin Zhang, Hao Pan). Senior authors with significant contributions (Carsen Stringer, Huiping Liu). Correspondence to: Huiping Liu, MD, PhD, Northwestern University, 303 E Superior St, Chicago, IL 60611. Carsen Stringer, PhD, HHMI Janelia Research Campus.

## Abstract

Circulating tumor cells (CTCs) and immune cells form dynamic multicellular ecosystems in blood, but their spatial organization and clinical relevance have not been systematically characterized. We developed the Cell and Cluster Identification Program (CCIP), an artificial intelligence–based framework that analyzes routine multiplex immunofluorescence blood scans to segment cells, identify CTCs and five immune lineages with high accuracy, and quantify multicellular clusters and tumor–immune interactions. Applying CCIP to 2,693 blood scans from 1,399 patients, we profiled over 60 million cells (>7 million multi-cell clusters) and linked imaging-derived features to patient outcomes. Correlated with circulating-tumor DNA mutation burdens, a 14-feature image model predicted overall survival in breast cancer, outperformed clinicopathologic variables and CTC enumeration, and generalized to prostate cancer. Prognostic imaging signatures were also associated with therapy response-related progression-free survival as well as with single-cell RNA sequencing–derived immune suppression states, connecting circulating tumor–immune architecture with systemic immune dysfunction.

## Introduction

Liquid biopsies offer a noninvasive window into tumor dissemination and host immunity, yet their clinical readouts remain strikingly reductionist. Circulating tumor cells (CTCs) rarely travel as solitary entities; instead, they form multicellular assemblies with other tumor cells and immune cells, structures that are associated with markedly enhanced metastatic potential and altered immune surveillance (*1–7*). While both homotypic and heterotypic CTC clusters have been implicated in metastasis, tumor plasticity, and immune evasion (*1–7*), these circulating architectures remain largely unexplored in clinical practice. Current liquid biopsy analyses typically collapse this complexity into simple enumeration (*1, 2, 7–9*), overlooking cellular morphology, spatial organization, and tumor–immune interactions that may encode critical biological and prognostic information.

Liquid biopsy imaging platforms are already widely deployed in clinical oncology, most prominently through FDA-cleared systems that enrich and image circulating cells using multiplex immunofluorescence (*8*). These platforms generate standardized, high-resolution images of tumor and immune cells across thousands of patients each year. However, downstream analysis remains largely manual and categorical (such as CTC enumeration) rather than interrogating multicellular architecture or spatial relationships. As a result, much of the information captured in routine blood cell imaging—particularly features reflecting tumor–immune organization—is underutilized in both biological investigation and clinical risk assessment.

Advances in artificial intelligence (AI) have enabled high-dimensional analysis of tissue images, uncovering previously inaccessible links between morphology and clinical outcomes (*10–14*). However, many of those lack associations with molecular signatures. Meanwhile, the rich imaging data already generated by standard liquid biopsy platforms have not been leveraged to interrogate circulating tumor–immune organization at scale. Here we present the Cell and Cluster Identification Program (CCIP), an AI-based imaging analytics framework that transforms routine multiplex immunofluorescence blood scans into quantitative profiles of circulating tumor and immune cells, their multicellular clusters, and spatial interactions. Based on a large dataset of blood cell imaging scans via CellSearch from patients with various cancers, CCIP integrates our open-source AI-based cell segmentation tool, Cellpose (*15–17*), for cellular, subcellular, and intercellular imaging feature extraction. By applying CCIP across thousands of clinical blood scans, we test whether circulating tumor–immune architectural states are reproducible, prognostic, and biologically informative, and whether they reflect functional immune states measured by single-cell transcriptomics.

By integrating population-scale liquid biopsy imaging with clinical outcomes and molecular immune states, this study seeks to establish circulating tumor–immune architecture as a biologically informative dimension of cancer progression and patient risk.

## Results

### Overview of CCIP

To establish a pilot AI-driven imaging analysis platform for automated identification of blood cell types, CTCs, and clusters, we leveraged a clinically acquired multiplexing CellSearch immunofluorescence imaging dataset from 2,693 blood scans of 1,358 patients with diverse cancers (mostly breast and prostate) and other diseases, spanning multiplexing channels for cytokeratin (CK), CD45, nuclear DNA (DAPI), and a customized marker (e.g., HER2) (**Fig. 1A, Supplementary Table 1**).

**Figure 1.**
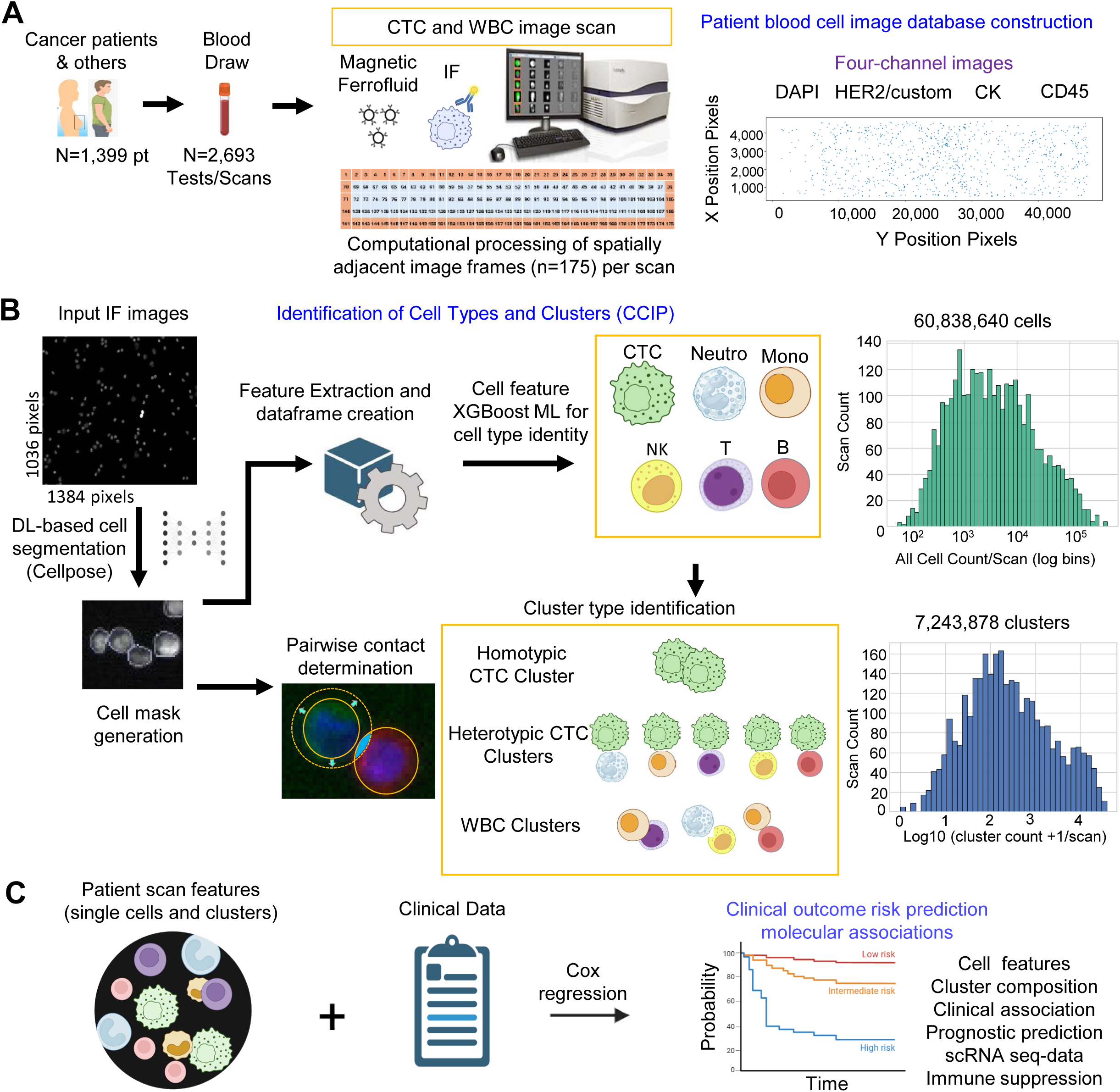
Overview of CCIP pipeline for image database construction, automated identification of cells and clusters, and clinical outcome prediction. **A.** Human blood cell image database construction, including biospecimen collection from patients with cancer and other diseases (N=1399 with 2,693 blood tests), CellSearch bead enrichment, four channel immunofluorescence staining, and whole slide (175-frame) image scanning. **B.** AI-enabled image processing, including Cellpose-based cell segmentation, imaging feature extraction, and XGBoost ML-based identification of cells and clusters (CCIP), detecting over 60 million cells (CTC, neutrophils, monocytes, T cells, B cells, and NK cells) and 7 million clusters (homotypic CTC clusters, heterotypic CTC-WBC clusters, and WBC clusters). The distribution histograms show scan number across the range of cell number (top panel) and cluster number (bottom panel) per scan, respectively, for all image scans. **C.** Patient blood test-derived image feature analyses for (1) prediction of clinical outcomes (overall survival) using a Cox regression proportion hazard model, and (2) association with molecular and cellular features such as immune suppression states (scores).

The CCIP pipeline was established to integrate multiple functions seamlessly: (1) Cellpose-based cell segmentation as the foundation for multi-channel fluorescence image feature extraction. Among four tested Cellpose models(*15–17*), the original model outperformed others (Denoise, Deblur, and Oneclick) with Pearson’s r = 0.976 in validations (**Fig. 1B, Supplementary Fig. S1A-E**) and was chosen to process our image dataset. (2) Automated identification of over 53 million cells and classification into CTCs (DAPI^+^CK^+^CD45^−^) and five manually non-distinguishable immune cells (DAPI^+^CK^−^CD45^+^), including neutrophils, monocytes, T cells, B cells, and NK cells. (3) Intercellular cluster identification, such as homotypic and heterotypic CTC clusters as well as diverse immune clusters. and (4) Cellular and cluster signature characterization, including image features, composition statistics, and intercellular contact strength (**Fig. 1B, Supplementary Table 1**). The CCIP-derived imaging features of the circulating tumor-immune ecosystem were incorporated into a Cox proportional hazards model to predict clinical outcomes (overall survival) and connect with functional immune cell states, such as the immune suppression score generated from our single-cell RNA sequencing data (**Fig. 1C**). The platform details and clinical impact of the CCIP are shown in the following sections.

### CCIP-driven identification of CTCs and immune cells

We hypothesized that an AI-powered imaging feature analysis platform will accelerate the lineage and spatial characterization of tumor-immune ecosystems for prognostic evaluation. To achieve machine learning-based cell identification of CTCs and various immune cells with limited immunofluorescence markers, we first isolated ground truth cells for CellSearch-loaded image scans, and trained the CCIP with 80% of the cells using imaging feature based XGBoost modeling, including immune cell lines (Jurkat and THP-1) and patient blood cells (neutrophils, monocytes, T, B, and NK cells). Then the remaining 20% of isolated cells and/or cells from independent datasets were mixed computationally to assess the performance of CCIP in cell identification (**Fig. 2**, **Supplementary Figs S2-S4**).

**Figure 2.**
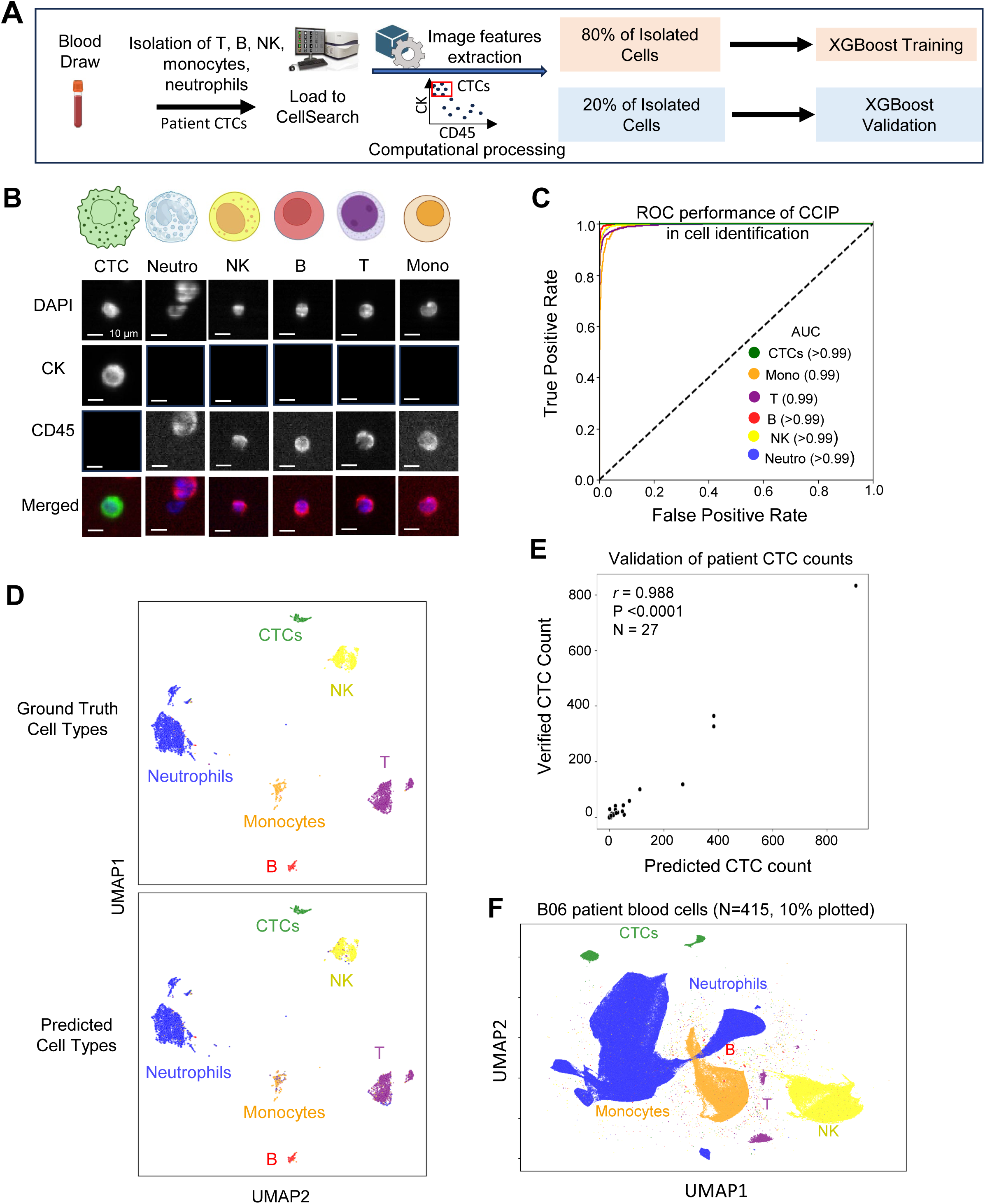
CCIP-driven identification of CTCs and WBCs. A. Schematic of ground truth blood cell isolation, including human WBCs (neutrophils, monocytes, T, B, and NK) and CTCs, using selection beads to generate IF images from Cellsearch for training (80% of data) and validation (20% of data) of CCIP classification model. B. Representative images of six cells types (CTC, monocyte, T, B, NK, and neutrophil) of flow sorted ground truth cells loaded to CellSearch platform for imaging, in three individual fluorescence channels, DAPI, CK, and CD45, and the merged channel. C. Receiver operating characteristic (ROC) curves illustrating CCIP model performance across 6 cell types (CTC, Neutrophils, T cells, NK cells, Monocytes, and B cells) with AUC >=0.99. D. Supervised UMAP projections comparing ground truth (top panel) and predicted (bottom panel) labels for the validation set of 8,576 blood cells (3,985 neutrophils, 2,006 T cells, 1,426 NK cells, 465 Monocytes, 263 B cells, 431 CTCs). E. Scatter plot comparing the verified report (ground truth) versus predicted CTC counts for individual cancer patient samples in the testing data (N=27). Each point represents a single sample. The Pearson’s correlation coefficient *r* and associated P value displayed. F. Supervised UMAP depicting 10% of total cells annotated from the CellSearch images of B06 breast cancer cohort (N=415) using CCIP platform, including 594,701 neutrophils, 153,953 NK cells, 123,448 monocytes, 28,733 CTCs, 17,008 T cells, and 2,424 B cells.

To establish a training set of CTCs, we computationally isolated high-confidence CTC candidates from CellSearch-loaded image scans. Specifically, image derived datapoints from five randomly selected patients were selected based on events that exhibited high cytokeratin signal and low CD45 signal, along with morphological features consistent with CTCs. Images of each CTC used for training were visually inspected confirming CTC identity. These verified CTCs were then incorporated into the CCIP training dataset, allowing for the ability to distinguish CTCs from other cell types (**Fig. 2A)**

From each cell, over 200 imaging features were extracted from cytoplasmic (CK and CD45) and nuclear (DAPI) channels into six categories for training: gradient (44), architecture (40), intensity (36), Fourier shape (32), cell morphology (20), and nuclear morphology (20) (**Supplementary Fig. S2A-C**). CCIP showed remarkable predictive power with its ability to not only recognize CTCs, but also distinguish morphologically similar immune cells (neutrophils, monocytes, T, B, and NK cells). The classification performance remained near perfect across all tested blood cell types, with the receiver operating characteristic curve (ROC) area under the curve (AUC) values approaching 0.99-1.00 (**Fig. 2A-C, Supplementary Fig. S3**). Supervised UMAP projections revealed strong alignment between predicted and ground truth labels (**Fig. 2D, Supplementary Fig. S2C-F**).

CCIP demonstrated high accuracy in automated reporting of CTC counts in patients, with a strong correlation (Pearson’s r = 0.988, p < 0.0001) between computationally predicted versus manually verified CTC counts in 27 randomly selected patients from the testing data (**Fig 2E**). When applying CCIP to blood image scans from the B06 cohort of breast cancer patients (n=415), six distinct cell types were grouped across patients in predicted label-guided UMAP projections (**Fig. 2F**). Feature importance analysis highlighted key discriminators such as cell size, signal intensity, and nuclear size (**Supplementary Fig. S2F-H, S4A-B**). Imaging features used for classification showed strong internal correlations within categories such as cell shape, fluorescence intensity, gradient, architecture, and nuclear morphology (**Supplementary Fig. S5**).

### CCIP characterizes spatial interactions and multicellular clusters with clinical utility

We next expanded the capabilities of the CCIP platform to specify spatial cell-cell interactions, such as the detection of multicellular clusters of CTCs and leukocytes. The CTC clusters are more likely to seed metastases than individual CTCs, and the detection of CTC clusters in cancer patients correlates with shorter survival, earlier relapses, and more aggressive disease symptoms compared to those with only single CTCs (*1–7, 18, 19*).

To enable automated spatial and intercellular cluster detection, CCIP incorporated a graph-based algorithm, comprising two key steps, (1) pairwise cellular interaction determination via optimized mask dilation (4 pixels) (**Supplementary Fig. S6A**), and (2) cluster characterization by identifying connected components within a cell graph (**Fig. 3A**). When integrated with cell type predictions, CCIP reliably identified intercellular clusters of any size and composition, including homotypic and heterotypic CTC clusters and diverse immune cell aggregates or clusters of various types (**Fig. 3B**). A validation study with 175 image frames demonstrated strong concordance between ground-truth (manually verified cluster counts) and CCIP-predicted cluster counts (Pearson coefficient r = 0.949, p < 0.0001) (**Fig. 3C, Supplementary Fig. S6B)**. Across all image scans, a positive correlation was also observed between cell counts and cluster counts (**Fig. 3D**).

**Figure 3.**
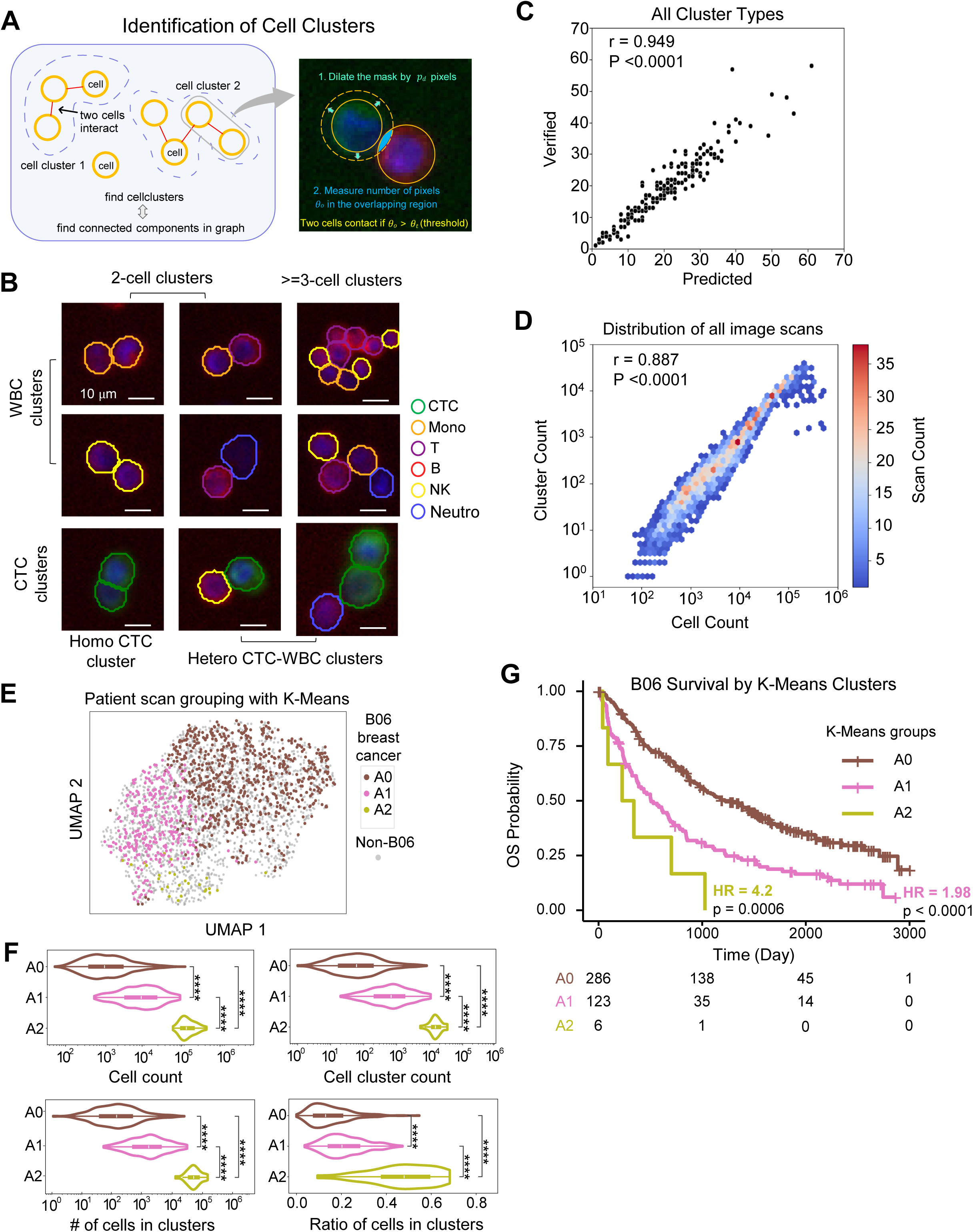
CCIP enables cell cluster detection and unsupervised patient stratification using abundancy features of cells and clusters. A. Schematic of the CCIP for cell cluster detection. The pairwise cell contact were first determined by the “Dilation-Intersection” procedure (*p_d_*-pixel steps for mask dilation; *θ_o_*-number of pixels in overlapping region; *θ_t_*-threshold for contact), and then contacting cells were aggregated into a complete cluster by searching connected components in a cell graph with individual cells as nodes and pairwise contact as edges. B. Representative image crops for immune cell (WBC) clusters, homotypic and heterotypic CTC clusters spanning from two-cell clusters to multi-cell (>2) clusters. Cell boundaries were annotated with colors indicating six cell types (CTCs, neutrophils, monocytes, T, B, and NK cells) using the CCIP pipeline. C. Scatter plot comparing computationally detected cell cluster numbers with manually verified counts for all cluster types (homo/hetero tumor clusters and WBC clusters). The Pearson’s correlation coefficient r and associated p value show strong linear correlation between two. D. Hexbin plot illustrating distribution of all image scans (blood tests) by cell count and cluster count. Each hexagonal bin represents a range of cell and cluster count values, and the color intensity reflects the number of scans falling within that range. E. UMAP of the spatial landscape for three cell/cluster abundance groups (A0, A1, A2) of patients with blood cell image scans, stratified by unsupervised K-Means clustering algorithm via the integrated abundancy features (full list see Supplementary Fig. 6c) about individual cells and cell clusters in each scan. A0-low abundancy, A1- medium abundancy, and A2-high abundancy, showing the B06 breast cancer patient cohort (n=1043 scans) in colors at foreground and all other cohorts in gray background (n=1650 scans). F. Violin plots showing distinct patterns of cell/cluster abundancy related features across three groups in B06 cohort, including total number of cells, total number of cell clusters, number of cells forming clusters, and ratio of cells forming clusters. The first three plots utilize log scale with base 10. Mann-Whitney U test was conducted across every two groups with star annotation (*: 0.01 < p <= 0.05; **: 0.001 < p <= 0.01; ***: 0.0001 < p <= 0.001; ****:p <= 0.0001). Refer to Supplementary Fig. 6d for distribution of three contact strength related features. G. The Kaplan-Meier curve showing overall survival probabilities for three abundancy groups (A0, A1, A2) of B06 breast cancer cohort, stratified by K-Means. Compared to A0, distinct survival patterns with HR, 95% CI, and log rank p-values for A1 (1.98, 1.55 to 2.53, p<0.0001) and A2 (4.20, 1.85 to 9.51, p=0.0006) demonstrate image feature-based patient stratification and outcome prediction in an unsupervised way.

We further profiled the spatial landscape of the cell and cluster features across patient-level blood cell image scans. Using unsupervised K-Means grouping on comprehensive cellular and cluster metrics, we stratified patient scans into three abundance groups: A0 - low abundance, A1 - medium abundance, and A2 - high abundance (**Supplementary Fig. S6C**). To quantify intercellular tumor–immune interactions, CCIP computed interaction strength between contacting cells using three metrics: minimum boundary distance, intersection-over-union ratio, and shared boundary length (**Supplementary Fig. S6D**). UMAP visualization revealed distinct A0, A1, and A2 groups, as shown in multiple cohorts of breast and prostate cancer patient scans (B06, PACE, and U08) (**Fig. 3E, Supplementary Fig. S7A-B**). Key abundance metrics—including total cell count, cluster count, number of clustered cells, and cluster-to-cell ratios—differed significantly across groups (**Fig. 3F, Supplementary Fig. S7C-D**).

Importantly, the abundance-based stratification (A0-A2) of the patient blood sample scans, performed without incorporating clinical or demographic data, revealed significant differences in overall survival, as shown by Kaplan-Meier blot analysis with hazard ratios (HR) of 1.98 (95% CI 1.55 to 2.53) and 4.2 (95% CI 1.85 to 9.51) for A1 and A2, respectively (log-rank test p < 0.0001) (**Fig. 3G**). These findings underscore the clinical utility of CCIP for unsupervised blood cell image analyses and prognostic stratification.

### CCIP image feature-derived risk score outperforms clinical pathological risk model

To identify CCIP -derived cellular and cluster image features associated with patient survival, we compiled quantitative metrics from blood cell images (n = 1,653 parameters), including cell counts, interaction scores, and averaged imaging features of various cell types and clusters. We then applied LASSO regularization(*20*) to select parameters potentially linked to patient outcomes, using the B06 patient cohort which was randomly divided into a training set (80% of the patients with n = 556 blood image scans) and a testing set (20% of the patients with n=152 scans).

With the training data, we constructed multivariate Cox proportional hazards models incorporating imaging features, disease stage, tumor grade, and patient age. The top list of CCIP-derived 14 imaging features (CCIP-14), involving CTCs, monocytes, NK, B, neutrophil, and T cells, were found to be significantly associated with survival (**Fig. 4A**). They are subsequently integrated with clinical characteristics, including age, stage, and grade (ASG) (**Fig. 4B**). We created a CCIP-14 image feature-derived model as well as a combinational model incorporating CCIP-14 image parameters along with ASG for risk analysis and outcome prediction.

**Figure 4:**
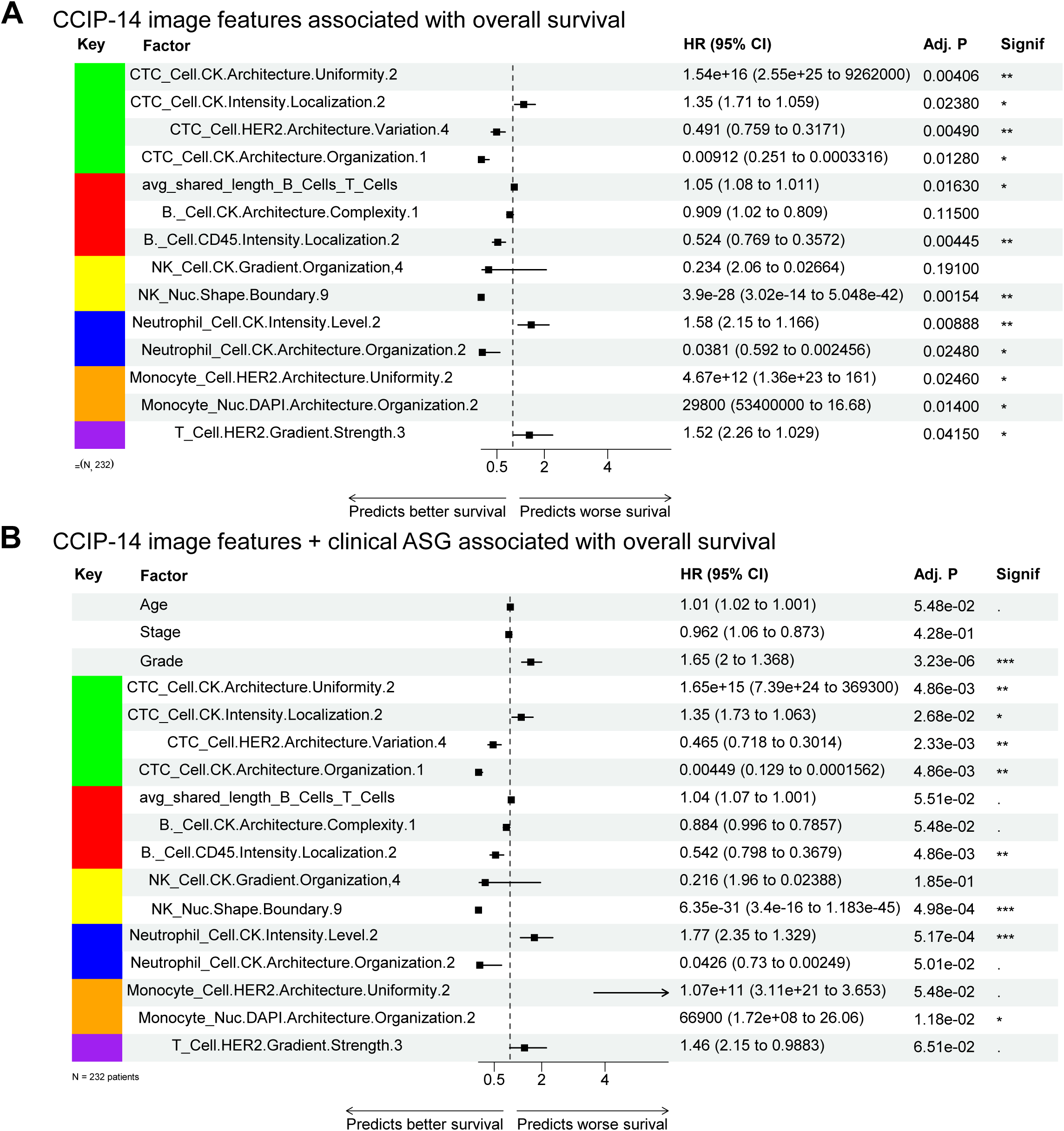
CCIP image features predicting patient overall survival. A. Top 14 CCIP image features of cells and clusters plotted with hazard ratios for patient overall survival, using a multi-variable Cox proportional hazards model. All cell and cluster image features were evaluated using LASSO regularization to identify the features most associated with patient overall survival. P values adjusted using FDR algorithm. B. CCIP-14 image features plus clinical and pathological features of age, stage and grade (ASG) in association with patient overall survival, analyzed using a multi-variable Cox proportional hazards model. P values adjusted using FDR algorithm.

A composite risk score based on the CCIP-14 imaging features alone was generated via the Cox Proportional Hazards model and stratified patients in the test set into high and low risk categories, with distinct survival probabilities (HR = 3.65, 95% CI 2.41 to 5.52, p<0.0001) (**Fig. 5A**). The predictive performance of the CCIP-14 risk score was evaluated using time-dependent ROC curves, yielding high sensitivity and specificity AUCs (*21*) (0.836 at 1 year) (**Fig. 5B**). In the meantime, the CCIP-14 plus ASG model has a lower hazard ratio (HR = 2.40, 95% CI 1.59 to 3.63, p<0.0001) (**Fig. 5C**), with comparable time-dependent ROC AUC values (0.853 at 1 year) (**Fig. 5D**). Notably, the clinical benchmark ASG-based risk scores in this cohort were not significantly associated with overall survival and did not stratify patients at all, showing low AUC performance (**Supplementary Fig. S8A-B,** HR=0.993, 95% CI 0.51 to 1.93, p=0.98), likely due to the cohort containing disproportionate patients at a high risk from their disease (**Supplementary Table 2**). Notably, the CCIP-14 image risk model also outperformed CTC enumeration (counts>=5, or <5) method, which an existing liquid biopsy model based on CellSearch reports (**Supplementary Fig. S8C-D,** HR=2.18, 95% CI 1.42 to 3.34, p=0.0003).

**Figure 5:**
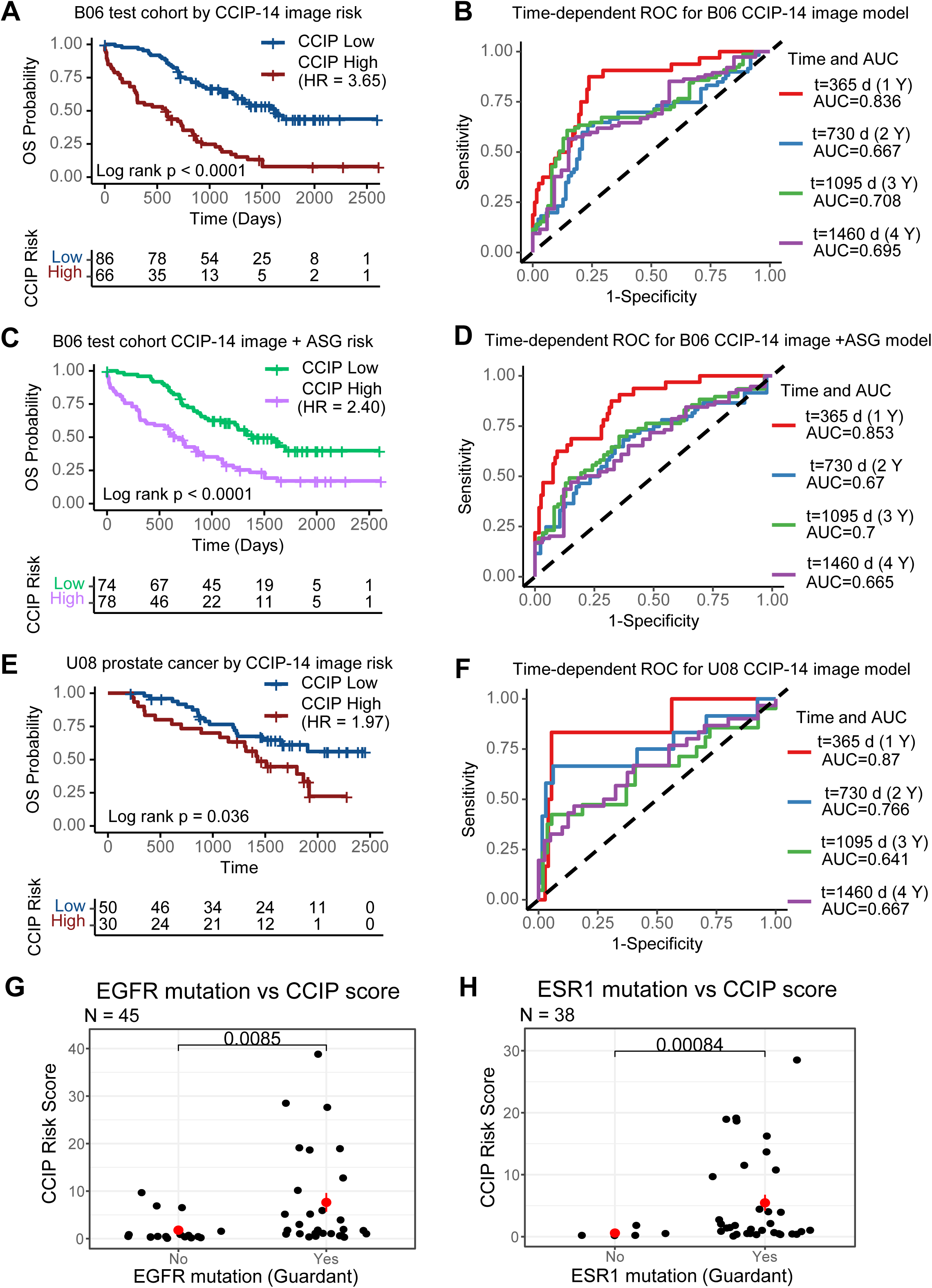
CCIP image-derived risk score outperforms clinicopathological features in prediction of patient overall survival. A. Kaplan-Meier overall survival curve on B06 breast cancer test data (20% of patients, n = 152 scans) with time from date of blood draw, using CCIP-14 image feature-derived risk score model for high or low risk stratifications. Cox proportional hazard model was developed using B06 training subset (80% of patients, n=608 scans), from which high-vs-low cutoffs were made. HR=3.65, 95% CI (2.41 to 5.52), log rank p < 0.0001. B. Time-dependent receiver operator characteristic (ROC) was calculated for the risk score generated in (a). ROC was calculated at 1, 2, 3 or 4 years (Y) after imaging. AUC was calculated for each time point. C. Kaplan-Meier survival curve using image + ASG (age, stage, and grade) model on B06 test data (n = 152 scans). Cox proportional hazard model was developed using B06 training subset, and high-vs-low cutoff levels were made based on that data. HR=2.40, 95% CI (1.59 to 3.63), log rank p < 0.0001. D. Time-dependent ROC was calculated for the risk score generated in (c). ROC was calculated at 1, 2, 3, and 4 years (Y) after imaging. AUC was calculated for each time point. E. Kaplan-Meier survival curve using images-only model on U08 (prostate cancer) validation data (n = 80 scans). Scores and stratifications were generated using the model based on B06 training data. HR=1.98, 95% CI (1.03 to 3.77), log rank p =0.036. F. Time-dependent ROC was calculated for the risk scores generated in (g). ROC was calculated at 1, 2, 3 and 4 years (Y) after imaging. AUC was calculated for each time point. G. Comparison of risk scores among patients with EGFR mutations as assayed by NGS vs patients without EGFR mutations H. Comparisons of risk scores among patients with ESR1 mutations as assayed by NGS vs patients without ESR1 mutations

In addition to the CCIP-14 image feature-derived risk scores, we also tested the CCIP top 40 image feature (CCIP-40) risk model (**Supplementary Fig. S9**). CCIP-40 feature scores stratified the B06 breast cancer cohort with high HRs, either by quartiles or by median cutoff (**Supplementary Fig. S8E-F**).

To generalize our CCIP models across cancer types, we expanded the CCIP risk scoring, including CCIP-14 and CCIP-40 image feature signatures derived from the breast cancer training set to an independent prostate cancer data set, NU18U08 (U08) (n=80). Stratified by the CCIP-14 and CCIP-40 image feature risk scores, the high-risk group with prostate cancer showed statistically significant HR=1.97 (95% CI 1.03 to 3.77, log rank p=0.036) for CCIP-14 and HR=2.14 (95% CI 1.12-4.11, log rank p=0.022) for CCIP-40 (**Fig. 5E, Supplementary Fig. S9G**). Similarly, the predictive performance of CCIP-14 for the independent U08 prostate cancer validation cohort achieved high AUC values of the time-dependent ROC curves (0.87 for 1 year) (**Fig. 5F**), demonstrating the generality of CCIP risk scores across multiple epithelial cancers.

A subset of NU16B06 patients have had next-generation sequencing (NGS) performed on circulating tumor DNA (ctDNA). To test whether CCIP scores were correlated with mutations identified by NGS, we compared CCIP scores from patients with or without mutations, as identified by Guardant ctDNA sequencing. Patients with an ESR1 mutation had a higher CCIP score than patients without such mutations (n = 45, p = 0.0085) (**Fig. 5G**). Likewise, patients with EGFR mutations had higher CCIP scores than patients without EGFR mutations (n = 38, p = 0.00084) (**Fig. 5H**). This suggests that CCIP scores are associated with downstream effects from underlying cancer-driving mutations.

### CCIP image features associated with progression-free survival (PFS) and immune suppression

To identify CCIP imaging features associated with therapy outcomes, we applied LASSO-regularized Cox proportional hazards regression to select features linked to chemotherapy progression-free survival (PFS) in a proof-of-concept study. We benchmarked six penalization strategies under 5-fold × 10-repeat cross-validation (**Supplementary Fig. S10A-B**). Among the penalized regression strategies evaluated by repeated cross-validation, the regular Lasso demonstrated both high time-dependent AUC at 90, 180, and 365 days and low cross-fold variability, indicating robust and stable out-of-sample discrimination. It was therefore selected for all subsequent analyses. Molecular subtype indicators (hormone receptor-positive or HR^+^, TNBC, or HER2^+^) and a stage III/IV binary indicator were also incorporated as unpenalized covariates, ensuring clinical context was retained regardless of regularization.

Before evaluating the CCIP imaging model, we first confirmed that our PFS endpoint — anchored to therapy initiation (t₀) with baseline imaging derived from the scan closest to but no later than t₀ within a 60-day window, and progression events defined from treatment response records — faithfully recapitulated established clinical prognostic hierarchies. The LOOCV-derived continuous risk score stratified the 107-patient chemotherapy cohort (40 progression events [38%]; median PFS 229 days) into significantly distinct survival groups. Dichotomizing patients at the median LOOCV risk score yielded a hazard ratio of 3.87 (p < 0.0001; **Fig. 6A**), and tertile stratification further resolved three prognostically distinct groups, with hazard ratios of 2.54 and 3.96 for the intermediate- and high-risk groups relative to low risk, respectively (p = 0.0034; **Fig. 6B**). The predictive performance of the risk score was evaluated using LOOCV-based time-dependent ROC curves, yielding consistent discrimination across clinically relevant horizons: AUC of 0.761 at 90 days, 0.734 at 180 days, 0.770 at 270 days, and 0.754 at 365 days (**Fig. 6C**).

**Figure 6.**
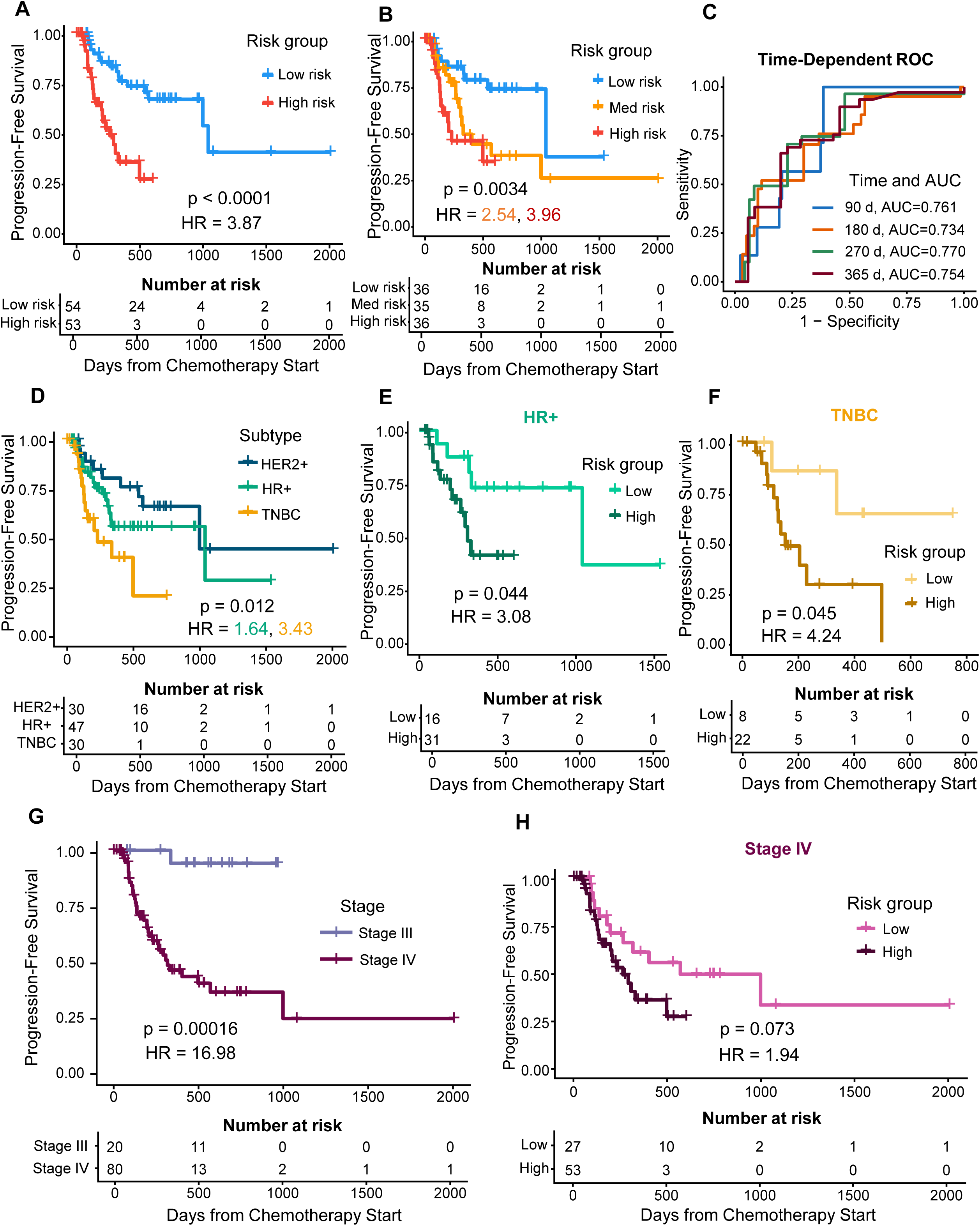
CTC imaging features associated with progression-free survival (PFS) in breast cancer patients receiving chemotherapy. A. Kaplan–Meier (KM) survival curves for PFS stratified by LOOCV CTC risk score dichotomized at the cohort median (Low risk, *N* = 54; High risk, *N* = 53). KM curves were estimated using the *survfit* function from the *survival* R package and compared between groups with the log-rank test. A leave-one-out cross-validated (LOOCV) composite CTC risk score was derived from LASSO-penalized Cox regression applied to baseline CTC imaging features. B. KM curves for PFS using tertile-based stratification of the LOOCV CTC risk score (Low, *N* = 36; Medium, *N* = 35; High, *N* = 36). C. LOOCV time-dependent receiver operating characteristic (ROC) curves evaluating the discriminative performance of the CTC composite score at 90, 180, 270, and 365 days from chemotherapy initiation. Area under the ROC curve (AUC) values are shown for each time point. All analyses were restricted to patients with complete baseline CTC scan data and available clinical outcome data. D. PFS stratified by molecular subtype: HER2-positive (HER2+, *N* = 30), hormone receptor-positive/HER2-negative (HR+, *N* = 47), and triple-negative breast cancer (TNBC, *n* = 30). Subtype was associated with significantly different PFS outcomes (log-rank *p* = 0.012), with HER2+ patients showing the most favorable prognosis and TNBC patients exhibiting the most rapid disease progression. E. KM curves for PFS within the HR+ molecular subtype (Low risk, *N* = 16; High risk, *N* = 31), stratified by the CTC risk score. F. KM curves within the TNBC subtype (Low risk, *N* = 8; High risk, *N* = 22). G. PFS stratified by disease stage at enrollment: Stage III (*N* = 20) and Stage IV (*N* = 80). Stage IV patients demonstrated markedly inferior PFS compared to Stage III patients (log-rank *p* = 0.00016), consistent with the known prognostic impact of metastatic disease. These distributions provide a clinical context for interpreting CTC-based risk stratification analyses presented in the main figures. H. KM curves within Stage IV patients (Low risk, *N* = 27; High risk, *N* = 53). Log-rank *p*-values are shown for all stratified comparisons.

We found the PFS differed significantly by molecular subtype (p = 0.012; **Fig. 6D**). TNBC patients showed the worst outcomes relative to the HER2^+^ subgroup (HR = 3.43), while HR+ patients exhibited intermediate PFS (HR = 1.64), largely consistent with the known chemotherapy response diversity across breast cancer subtypes. Within each molecular subtype, the CCIP imaging-derived risk score retained independent prognostic value. Among the HR^+^ patients (N = 47), high-risk individuals experienced a 3.08-fold greater hazard of progression than low-risk patients (p = 0.044; **Fig. 6E**), and in TNBC patients (N = 30), the corresponding HR was 4.24 (p = 0.045; **Fig. 6F**).

Notably, the B06 patients with Stage IV disease (N = 80) experienced markedly worse PFS than those with Stage III (N = 20; HR = 16.98, p = 0.00016; **Fig. 6G**). Among Stage IV patients alone, the LOOCV risk score further suggest potential prognostic stratification beyond stage classification (HR = 1.94, p = 0.073; **Fig. 6H**), suggesting that CCIP imaging features capture residual survival heterogeneity not likely explained by clinical staging. Taken together, these findings demonstrate that baseline CCIP imaging features provide robust, therapy-contextualized prognostic stratification in breast cancer patients receiving chemotherapy, with consistent performance across molecular subtypes and disease stages.

To link CCIP imaging-derived survival factors with molecular pathways of systemic immune cell states, we analyzed single-cell RNA sequencing data with WBCs (immune cells) collected from breast cancer patients (N=18 from the B06 cohort) (**Supplementary Fig. S11A**). We curated an immune suppression–related gene signature, spanning immunoregulatory cytokines and pro-angiogenic factors (*22*), metabolic and enzymatic mediators of immune inhibition (*23*), and inhibitory immune checkpoint molecules associated with T cell exhaustion and functional impairment (*24, 25*). In addition, we included markers of alternatively activated or regulatory myeloid polarization (*24*), transcriptional programs characteristic of regulatory T cells (FOXP3) (*26*), key modulators of anti-inflammatory signaling and feedback control (*27, 28*), and the pro-tumorigenic inflammatory mediator SPP1(*29*). At the patient level, aggregation of single-cell immune suppression scores revealed substantial inter-patient variability in systemic immune states (**Supplementary Fig. S11B-C**).

To determine whether imaging-derived features indicate these molecular immune states, we next evaluated the association between survival and therapy response-associated imaging features and the scRNA-seq–derived immune suppression score. Univariate regression identified multiple imaging features significantly associated with immune suppression, spanning diverse cell types including monocytes, NK, neutrophils, and T cells (**Supplementary Fig. S10D-J**). Together, these findings indicate that CCIP imaging-derived features correlate with biologically meaningful variation in the tumor–immune microenvironment in addition to their association with therapy response.

## Discussion

This study demonstrates that imaging features derived from blood cells can serve as powerful prognostic biomarkers for predicting cancer outcomes, such as patient survival, and outperform the existing clinical stratification approaches based on patient age, stage, and grade as well as CTC enumeration. We have developed the CCIP pipeline, which integrates AI-based Cellpose segmentation (*15–17*) with ML-driven feature extraction to automate high-throughput image processing, cell detection, classification, and spatial characterization of CTCs and immune cells (neutrophils, monocytes, NK cells, T cells, and B cells). Beyond CTC enumeration, CCIP enables scalable analysis of multi-channel immunofluorescence signals at single-cell and intercellular levels to characterize spatial interactions and multicellular clusters. The CCIP pipeline can be feasibly applied to other imaging technologies for the detection and characterization of tumor cells and immune cells from various biopsies, using different imaging technologies (*3–5, 8, 9, 30, 31*).

Our findings support the emerging view that CTC-immune cell interactions serve as a critical determinant of metastasis and patient outcome, consistent with previous work showing that heterotypic CTC-leukocyte clusters enhance metastatic seeding efficiency compared with single CTCs(*18, 19, 32*). The ability of CCIP to capture these interactions at single-cell resolution allows for quantitative assessment of cellular spatial networks, previously unobservable with traditional enumeration-based assays.

As proof-of-concept, CCIP expands the capabilities of liquid biopsy imaging platforms to comprehensive profiling of tumor-immune ecosystems and survival risk stratification. Many of the patient survival-related imaging features are associated with immune suppression, implicating that the reciprocal modulation and dynamic interplay between tumor and immune cells impact cancer outcomes. The strong concordance between imaging-derived immune features and transcriptomic immune suppression scores from single-cell RNA sequencing further validates the biological relevance of the imaging-derived signatures, echoing recent studies linking morphological heterogeneity to functional immune phenotypes (*33, 34*).

Consistently, AI-driven digital imaging, such as digital pathology (*10–14*), mammography (*13, 35*), and other radiologic imaging (*36*), has transformed clinical diagnostic practice and enabled imaging-based prognostic modeling across multiple disease contexts. High-dimensional imaging features derived from these modalities have been shown to correlate with molecular programs and clinical outcomes (*10, 13*).

In summary, our study demonstrates the transformative potential of deep learning and machine learning in accelerating and enabling large-scale imaging data analyses of tumor immune ecosystem in blood biopsies and can readily extend the method to tissue images, as well as integrating imaging data with other modality of patient data such as digital pathology and spatial omics. By automating and enhancing the analysis of CTC-immune cell interactions, we present a robust framework for developing predictive models with direct clinical relevance. This work opens avenues for personalized treatment strategies, underscoring the impact of AI in advancing precision oncology.

## Materials and Methods

### Human subjects

All human subject research was performed under protocols approved by the Institutional Review Boards (IRBs) of participating institutions, with informed consent obtained from all patients or their legally authorized representatives. De-identified clinical and imaging data were analyzed to ensure the protection of participant privacy and confidentiality. The blood collection from patients has been approved by the Northwestern University IRB protocols STU00203283 (NU16B06 breast cancer), DF/HCC 17-101 (PACE breast cancer, NCT03147287), STU00207694 (NU18U08 prostate cancer), STU00214936, STU00203216, and STU00216869 for other cohorts and in association with clinical trials NCT03838367 and NCT04354701. All patients signed informed consent to be enrolled in this study.

Blood was collected in CellSave tubes (containing cellular fixative) and uploaded to the CellSearch® platform (Menarini Silicon Biosystems) for CTC analysis at the CTC Core Facility at Northwestern University. Briefly, approximately 7.5 mL of whole blood was collected from the patients into a 10-mL CellSave Preservative tube containing a cellular fixative. Blood specimens were maintained at room temperature (RT) and processed on Celltracks Autoprep® System with the special kits listed in the Supplementary Table 1. After the processing, the cartridges containing enriched cells were scanned and analyzed on Celltracks Analyzer II® System for manual and/or automated computational analyses. Positive tumor cells and interacted immune cells were claimed on the staining of fluorescently labeled antibodies against cytokeratin (CK), CD45, HER2, and with DAPI, et al. We specifically analyzed the three variables of interest, the presence of ≥ 5 CTCs/7.5 mL blood, HER2 + CTCs (or ER+, PDL1+, et al), and/or CTC homogeneous/heterogeneous clusters.

### Immune cell isolation

Peripheral blood mononuclear cells (PBMCs), polymorphonuclear leukocytes (PMNs) and red blood cells were separated by gradient centrifugation using Polymorphprep (Serumwerk Bernburg Product # 1895). PBMC sub-populations were isolated using EasySep human T cell isolation kit (StemCell Technologies product #17951), EasySep human NK cell isolation kit (StemCell Technologies product #17955), EasySep human monocyte isolation kit (StemCell Technologies product #19359), and EasySep human B cell isolation kit (StemCell Technologies product #17954). Isolated cell populations were added to pelleted red blood cells (RBCs) to create artificial samples containing only a single sub-population, and these artificial samples were run on the CellSearch platform. THP-1 cells (ATCC TIB-202) and Jurkat cells (ATCC TIB-152) were similarly combined with RBCs to create artificial samples.

### Flow cytometry analysis

The purity of immune cells isolated by the methods described above was evaluated using flow cytometry analyses. Briefly, Cells were then stained with the following antibodies: for T-cells and neutrophils, anti-CD45 Pe/Cy7 (BD Biosciences, #557748), anti-CD3 FITC (BD Bioscienes, #555339), anti-CD16 BV605 (Biolegend, #302039), anti-CD11b BV510 (Biolegend #301333), anti-CD66b PE/dazzle (Biolegend, #305121), anti-CD19 Percp/Cy5 (Biolegend, #302229), anti-CD56 AF700 (Biolegend, #318316) were used. For B-cells, monocytes, and NK cells, the following antibodies were used: anti-CD45 APC (BD Biosciences, #555485), anti-CD3 PE (BD Biosciences, #555340), anti-CD16 APC/Cy7 (Biolegend, #302017), anti-CD14 Percp/Cy5 (Biolegend, #325622), anti-CD11b PE/dazzle (Biolegend, #301347), anti-CD19 PE/Cy7 (Biolegend, #302215), anti-CD56 AF700 (Biolegend, #318316). Cells were stained for 30 minutes at 4 °C, then washed in 1x PBS and resuspended in 1x PBS containing 2% FBS and DAPI to mark dead cells and analyzed using the BD FACSymphony A5 SE Cell Analyzer. Flowcytometry data were analyzed using FlowJo v10.6.2.

### Cell segmentation

Cells from CellSearch immunofluorescence image scans, spanning four channels across 175 frames per scan (Supplementary Fig. 1), were segmented using Cellpose(*15–17*), an advanced deep learning-based tool designed for robust and accurate cell segmentation across diverse imaging modalities. Fluorescent images of cells were processed through our analysis pipeline, where a combination of models was applied for nuclear (Cellpose-Cytotorch_0) and cytoplasmic (Cellpose-TN1) segmentation respectively, which utilized DAPI for nuclear detection and cytokeratin/CD45 markers for cytoplasmic boundaries.

To evaluate segmentation performance, we compared cell counts predicted by the Cellpose model to the ground-truth counts under four distinct preprocessing conditions—Original, Denoise, Deblur, and Oneclick—with manually annotated counts. The Original pipeline yielded the strongest linear correlation (Pearson’s r = 0.976), with a fitted relationship of y = 0.88x + 4.23, outperforming the other models, and thus was used for subsequent analyses (Supplementary Fig. 1c). Cells were segmented at three levels using different channel combinations (Supplementary Fig. 1d): nucleus (DAPI), cytoplasm (CK, CD45), and whole-cell signals (DAPI, CK, CD45), accounting for heterogeneous or patchy staining patterns due to non-uniform marker distribution. Further quantitative evaluations of the Original pipeline across multiple Intersection-over-Union (IoU) thresholds demonstrated high precision, sensitivity, and positive predictive value (Supplementary Fig. 1e).

Segmentation masks generated by Cellpose were subsequently used for downstream quantitative analyses, including extraction of 257 (207 with customized HER2 channel features excluded) cellular features quantifying morphology, appearance and intensity of cells. These segmented masks provided a reliable foundation for further classification.

### CCIP-based cell type prediction

The CCIP model was established for the **C**ell and **C**luster **I**dentification **P**rogram, employing Extreme Gradient Boosting (XGBoost)(*37*) classifier for cell type prediction. This type of model utilizes and combines the predictions of multiple decision trees as well as uses gradient boosting framework, focusing on incorrect predictions, making it robust and accurate. Prior to training, 257 (plus customized channel) quantitative features were extracted from segmented cells and generated cell masks. These features included fluorescent intensity (36), morphological descriptors (40), Fourier shape (32) descriptors, and architecture-based measurements, including gradient (44) and architecture features (40). The customizable and interchangeable channel of 50 features were excluded from the training due to inconsistencies in the marker through patient scans. A dataset containing 207 features from 2,324 monocytes, 10,030 T cells, 19,924 neutrophils, 1,314 B cells, 7,133 NK cells, and 2,155 circulating tumor cells (CTC) were extracted and split 80:20 for training data and test data, respectively. The specified hyperparameters of the XGBoost model are as follows, ‘learning_rate’: 0.1, ‘max_depth’: 5, ‘n_estimators’: 100.

### CCIP-based cell cluster detection

Cell cluster detection in CCIP was performed using a two-step graph-based algorithm designed to capture spatially associated cells with high precision. In the first step, pairwise cellular interactions were defined through mask dilation with optimized range *p*_d_, in which each segmented cell mask was expanded outwards by four pixels to account for local cell–cell proximity (Supplementary Fig. 6a). We measured number of pixels *θ*_o_ in the overlapping region and compared it to a pre-defined threshold *θ*_t_ = 10. Adjacency relationships were then established between any two cells with *θ*_o_ greater than the threshold, forming an undirected cell graph. In the second step, clusters were delineated by identifying connected components (*38*) within this graph (Fig. 3a), enabling automated expansion from individual pairwise interactions to multi-cellular assemblies. When integrated with cell type annotations from the classification model, this approach allowed CCIP to systematically identify cellular clusters of arbitrary size and composition, including both homotypic and heterotypic circulating tumor cell (CTC) clusters as well as diverse immune cell aggregates.

### Model evaluation

Model performance was assessed on the test set using multiple classification metrics summarized in the table below. We plotted multiclass receiver operating characteristic curve (ROC) and calculated area under the curve (AUC). Feature importance scores were extracted from the trained model to identify the variables with the most predictive influence.

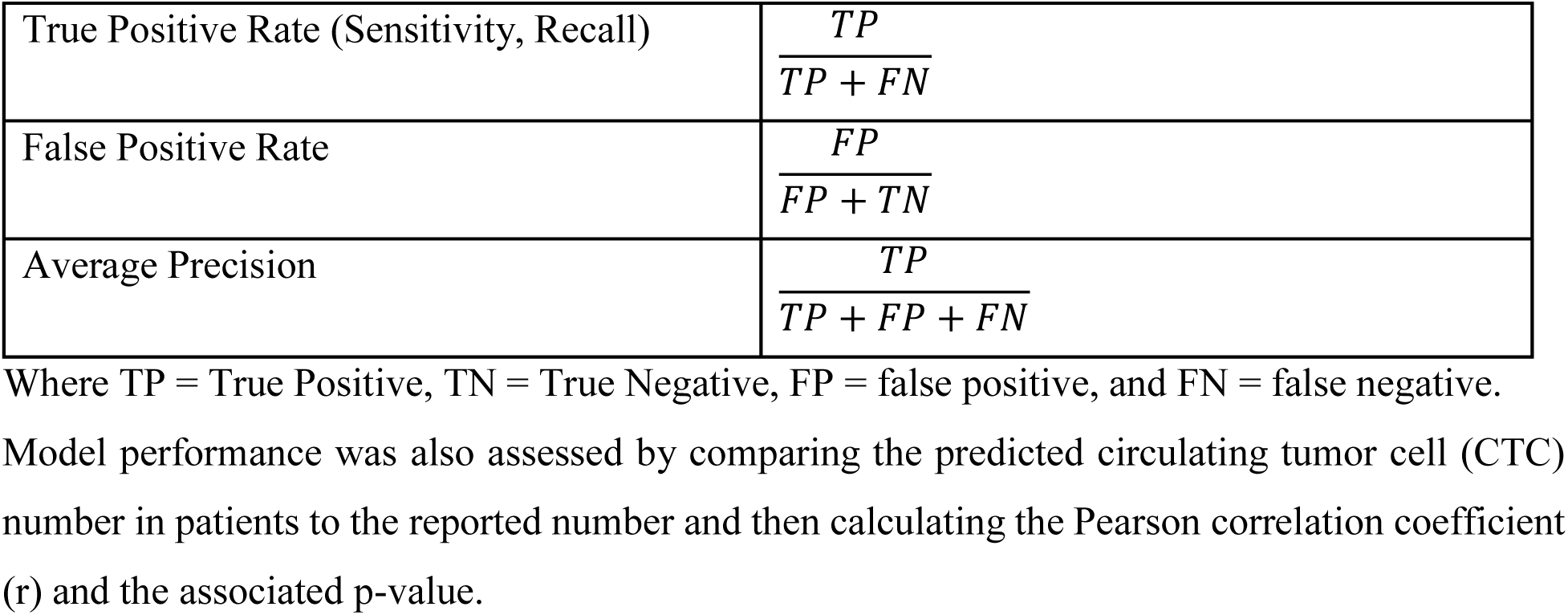

### Identification of imaging features associated with patient survival

Cell identity and cell cluster information were determined as in Fig 2 and Fig 3. For each CellSearch cartridge, cells were grouped based on identity and cell information across the 257 quantitative features were averaged for each cell type. These averaged cell data were combined with cluster data and cell counts, and cartridge data was filtered for the most recently collected sample for each patient in the NU16B06 cohort for a total of 1,653 variables.

The glmnet (*39–41*) implementation of LASSO regularization was used at 100 iterations to identify variables that could be associated with survival among NU16B06 patients. A total of 43 variables were identified using this method. Patients were dichotomized into high and low-risk groups based on median risk scores derived from imaging features. Log-rank tests were applied for four quartile groups in Kaplan-Meier plot. R package survival and survminer were used for survival analyses.

### Creation of risk scores for overall survival prediction

The NU16B06 cohort was divided into a training group (80% of patients) and a testing group (20% of patients). The training group was used to create a Cox proportional hazards model (*42*) and the underlying characteristics were evaluated for significance. Characteristics from that model that were significantly associated with survival were used to create a revised, 14-variable model which was then used to determine a risk score for the test group. Risk scores (*43*) were stratified into two groups using the training data and the stratification of the test data was plotted against survival in Figure 5 and Extended Figure 9. A baseline risk model was created using pathologist-determined grade and presence or absence of IHC staining for ER, PR or HER2. CTC counts were evaluated based on the CellSearch platform’s software. HR was calculated using a Cox proportional hazards test. P-value was calculated using Wald test and adjusted using the FDR method. Time-dependent receiver operating characteristic scores were created using risk scores at t=365 days (one year), t=730 days (two years), t=1095 days (three years), and t=1470 days (four years) and plotted in Fig 5(*44*). All analysis performed in R version 4.2.1(*45, 46*).

To create a baseline score, we used NU16B06 patients’ clinical data including cancer stage, tumor grade and patient age.

Prostate cancer (NU18U08) patients were analyzed in the same manner as NU16B06 patients, using the Cox proportional hazards model created for NU16B06 to predict a risk score.

### Therapy response-related progression-free survival association analysis

For chemotherapy, time-to-event was defined relative to therapy initiation (t₀). Baseline CTC imaging features were extracted from the scan closest to — and no later than — t₀, requiring the scan to fall within 60 days prior to therapy start (t₀ − t_image ∈ [0, 60] days). Progression events were defined as the first treatment record with best response of progressive disease (PD) occurring after t₀; patients without a recorded PD were censored at the last observed date, defined as the maximum of all scan and treatment record dates.

Kaplan–Meier curves (*47*) were used to estimate PFS distributions; groups were compared with the log-rank test. Univariate and subgroup hazard ratios (HRs) with 95% confidence intervals (CIs) were derived from Cox proportional hazards models (*48*).

A Lasso-penalized Cox regression model (*49, 50*) was trained on baseline CTC imaging features to predict PFS. Molecular subtype indicators (HER2+, TNBC, HR+) and a stage IV binary indicator were included as unpenalized covariates in all models. Model performance was estimated by leave-one-out cross-validation (LOOCV): for each patient, the model was trained on the remaining N−1 patients with the regularization parameter (λ) selected by up to 10-fold cross-validated deviance, and a linear predictor (risk score) was generated for the held-out patient. (*39, 51*) Patients were stratified into binary (median split) or tertile (33rd/67th percentile) risk groups from these LOOCV scores. Time-dependent discrimination was assessed by the cause-specific cumulative/dynamic AUC at 90, 180, 270, and 365 days using the timeROC (*21*) package, with 95% CIs derived from the influence function. Subgroup analyses by molecular subtype (HR+, HER2+, TNBC) and disease stage (III vs. IV) were performed independently within each stratum. All analyses were conducted in R (*45, 46*).

### Immunosuppressive score calculation

To evaluate the immunosuppressive potential of individual patients in our scRNA-seq dataset, we calculated an immunosuppressive score based on the expression of a panel of immunosuppressive genes, including *IL10, TGFB1, VEGFA, ARG1, IDO1, PDCD1* (PD-1), *CD274* (PD-L1), *CTLA4, LAG3, HAVCR2* (TIM-3), *NOS2, PTGS2* (COX-2), *CD163, MRC1* (CD206), *MSR1, FOXP3, IL4R, IL1RN, SOCS1, SOCS3,* and *SPP1*. Normalized expression values were extracted from the Seurat object, and the AddModuleScore function in Seurat (*52*) was used to calculate a gene module score for each cell. The mean immunosuppressive score across all cells from each patient was used to represent the patient-level (overall) immunosuppressive score, which was then used for downstream correlation analyses.

We curated an immune suppression–related gene signature, spanning immunoregulatory cytokines and pro-angiogenic factors (IL10, TGFB1, VEGFA) (*22*), metabolic and enzymatic mediators of immune inhibition (ARG1, IDO1, NOS2, PTGS2/COX-2) (*23*), and inhibitory immune checkpoint molecules associated with T cell exhaustion and functional impairment (PDCD1/PD-1, CD274/PD-L1, CTLA4, LAG3, HAVCR2/TIM-3) (*24, 25*). In addition, we included markers of alternatively activated or regulatory myeloid polarization (CD163, MRC1/CD206, MSR1)(*24*), transcriptional programs characteristic of regulatory T cells (FOXP3)(*26*), key modulators of anti-inflammatory signaling and feedback control (IL4R, IL1RN, SOCS1, SOCS3) (*27, 28*), and the pro-tumorigenic inflammatory mediator SPP1 (*29*).

The immune suppression score assesses a comprehensive panel of canonical immunosuppressive genes representing mechanistically interconnected regulatory axes, such as cytokine-mediated immune dampening, checkpoint-driven adaptive immune suppression, suppressive myeloid reprogramming, metabolic restriction of effector responses, and regulatory feedback circuits that reinforce immune tolerance. By integrating these convergent immunosuppressive programs into a unified module score, our immune suppression score can provide a quantitative, systems-level measure of immunoregulatory burden, enabling robust inter-patient comparisons and downstream correlation analyses with cellular composition, functional states, and clinical parameters.

### Image feature association with immune suppression scores

We analyzed patient-level blood-cell image–derived features in relation to immunosuppression scores across immune cell subsets (B, T, NK, monocyte, neutrophil) as well as an overall score generated from the single-cell RNA sequencing data. Features included morphological, textural, and intensity metrics derived from CK, HER2, CD45, and DAPI channels. To address missing data, we applied multiple imputation using the mice package (*53*), with CART for features and predictive mean matching for scores, selecting one completed dataset from five imputations. All features identified earlier (survival-associated and therapy response-associated) were then individually tested using univariable ordinary least squares (OLS) regression models, where each score was regressed on a single feature. From these models, we extracted β coefficients, 95% confidence intervals, and P-values. Visualization was performed using the forestplot (*54*) package, displaying effect sizes, confidence intervals, and P-values. All analyses were conducted in R (*45*).

### Association of CCIP score with NGS-identified mutations

Commercially-available NGS from Guardant was performed for NU16B06 patients as part of their care regimens at Northwestern Memorial Hospital. Patients with identified amplifications, deletions or missense mutations for EGFR or ESR1 were categorized as patients with mutations, while patients without these were categorized as patients without mutations. Scores were derived from the CCIP-14 model and compared by student’s T-test.

### Statistical Analysis

For all tests, p < 0.05 was considered significant. We reported Pearson’s correlation coefficient r and associated p-value for estimating linear correlation between manual count of instants and predicted numbers. The Mann-Whitney U test was applied to compare abundancy related features across three patient groups. Log-rank test was applied for four quartile groups in Kaplan-Meier plot. P values were adjusted when noted using the FDR method (also known as the Benjamini-Hochberg method). All graphs represent mean ± standard deviation unless otherwise specified.

## Acknowledgements

This work has been supported by the core facilities at Northwestern University including the Robert H. Lurie Comprehensive Cancer Center CTC Core, the Flow Cytometry Core, the Pathology Core, and the Quantitative Data Sciences Core. We are deeply grateful to the clinical team leaders and investigators Drs. Jonathan Strauss, Sarika Jain, Jindan Yu, Nausheen Akhter, and Seema Singhal as well as the patients who consented to donate their blood biospecimens. We thank all team members in the Liu Cluster and collaborative laboratories of Drs. Deyu Fang and Yuan Luo at Northwestern University and Dr. Massimo Cristofanilli’s laboratory at Cornell University.

## Funding

National Institute of Health/National Cancer Institute grant R01CA245699 (HL)

National Institute of Health/National Cancer Institute grant R01AI167272 (HL)

This project has been partially funded by NIH/NCI grant R01CA298232 (HL)

This project has been partially funded by NIH/NCI fellowship T32 CA009560 (DS)

Chan Zuckerberg Biohub Chicago (HL, LC)

American Cancer Society CSCC-Team-23-980420-01-CSCC (HL)

Prostate Cancer Foundation Challenge Award 2017CHAL2008 (JY, MH)

NIDDK training grant U2CDK129917 (AS)

NIDDK training grant TL1DK132769 (AS)

Lynn Sage Breast Cancer Research Foundation (HL)

Northwestern University Pharmacology start-up grant (HL)

## Author Contributions

Conceptualization: JRS, YS, ADH, YZ, HL, CS

Methodology: JRS, YS, ADH, YZ, LC, LZ

Investigation: JRS, YS, ADH, YZ, HP, FT, YH, DS, HA, AM, AS, JZ, HD

Supervision: HL, CS, DF, LZ, LC, MC, WJG, YL, MH, JY, LCP

J.R.S., Y.S., A.D.H, C.S., and H.L. conceived the idea presented here and wrote the manuscript. Y.Z. collected and provided all the raw image dataset derived from CellSearch platform with blood specimens of patients with cancer and other diseases. J.R.S. and Y.S. created CCIP for image feature analysis. A.D.H. identified the clinical outcome-related image features. H.P., F.T., Y.H. analyzed the association of survival-related features with single-cell RNA seq derived immune suppression. D.S., M.G., A. M., A.S., J.Z., H.D., and C.M. provided technical support and help in flow cytometry, sorting, cell isolation, and computational program development. Senior authors H.L., C.S., D.F., L.Z., L.C., M.C., W.J.G., Y.L., M.H., J.Y., and L.C.P. supervised experimental planning and computational implementation.

## Inclusion & Ethics Statement

This study was conducted in accordance with the highest standards of research integrity, ethical oversight, and inclusive scientific practice. We are committed to promoting equity, inclusivity, and representation in cancer research. The patient cohorts analyzed in this study reflect the diversity of individuals receiving clinical care at the participating institutions, and no exclusions were made on the basis of sex, gender identity, race, ethnicity, socioeconomic status, or other demographic factors unless dictated by clinical eligibility criteria. Where demographic or clinical variables were available, they were considered in downstream analyses to identify potential disparities and avoid the development of biased models.

All computational methods were developed and evaluated with attention to fairness, transparency, and reproducibility. The AI models used in this work were trained and validated on large, heterogeneous datasets to minimize bias and maximize generalizability across cancer types and patient subgroups. Model outputs were systematically assessed for consistency across demographic strata, and care was taken to avoid the introduction of algorithmic bias during model training or interpretation.

All authors affirm adherence to ethical standards for authorship, data handling, and reporting. No fabrication, falsification, or inappropriate data manipulation was performed. The study complies with institutional, national, and international guidelines on the responsible conduct of research and the ethical use of human data.

## Data availability

The data supporting the findings of this study are available in the Supporting Data Values file. Human scRNA-seq datasets generated in this study are available at GSE294399(*19*): https://www.ncbi.nlm.nih.gov/geo/query/acc.cgi?acc=GSE294399. The entire set of scanned immuno-fluorescence images of patient blood and pre-calculated image features and type prediction for individual cells, as well as cell-cluster features can be downloaded through Globus upon requests: https://app.globus.org/file-manager?origin_id=75af3eb6-b480-4f19-882c-ed079225b0ec&origin_path=%2F.

## Code availability

The code is available for testing using the supplementary package “CCIP_Code_For_test.zip” and upon collaboration request at https://github.com/Stephen2526/CTC_Image. Cell segmentation was conducted with Cellpose v3.1.0. Cellular features were calculated using HistomicsTK (v1.4.0, https://github.com/DigitalSlideArchive/HistomicsTK) and scikit-image (v0.24.0). Machine learning models for cell type identification and patient clustering were trained with xgboost (v3.0.0) and scikit-learn (v1.5.2). Python (v3.9.15), scipy (v1.13.1), pandas (v2.2.3), matplotlib (v3.9.4) and seaborn (v0.13.2) were used for data analysis and plotting figures. A series of R packages were used for patient survival analysis and association study with immunosuppression scores: R (v.4.2.1), Seurat (v.4.4.2), dplyr (v.1.1.4), tidyverse (v2.0.0), ggplot2 (v3.5.1), survminer (v0.4.9), enrichR (v3.3), dittoSeq (v1.16.0), EnhancedVolcano (v1.27.0), forestplot (v3.1.7), and survival (v3.7.0). FlowJo software (version 10.10.0) was used to generate flow cytometry plots and related data analysis.

