## Supplementary Tables 1-2, Supplementary Figures S1-S11 for "AI-enabled Spatial Profiling of Circulating Tumor-Immune Ecosystems Predicts Patient Outcomes Across Cancers"

#### Supplementary Figures S1-S11

### Supplementary Table 1. List of cancer and viral research projects

| Disease | Breast Cancer | | | | | | Prostate Cancer | Multiple Myeloma | COVID-19 | Total |
|---|---|---|---|---|---|---|---|---|---|---|
| Project | NU16B06 | NU20B02 | PACE | CD07_TNBC | NU16B09 | others | NU18U08 | CMMC | CEC_COVID | |
| Patient Number | 478 | 27 | 221 | 30 | 28 | 431 | 41 | 67 | 76 | 1,399 |
| Blood Test Number | 1,151 | 27 | 436 | 136 | 28 | 431 | 85 | 252 | 147 | 2,693 |
| Menarini Kit info | CTC | CTC | CXC | CXC | CXC | CTC and CXC | CXC | CMMC | CEC | |
| Customized Marker | HER2-FITC | N/A | ER-PE | CCR5-PE | PDL1-PE | HER2-FITC, SEMA4A, RANK, PODXL, PLXNB2, ICAM1, CD44, CD4, CD3, CD25, CD68, SNA | AR-PE | N/A | ACE2-FITC | |
| Total Cell Count | 18,067,330 | 24,528 | 21,559,358 | 5,172,697 | | 11,949,937 | 483,995 | 2,183,447 | 1,397,348 | 60,838,640 |
| CTC count | 574,518 | 108 | 177195 | 13,776 | | 329,545 | 10,429 | 907,358 | 20,477 | 2,033,406 |
| Monocyte Count | 2,475,316 | 738 | 2,818,124 | 336,328 | | 1,687,796 | 16,960 | 105,157 | 49,020 | 7,489,439 |
| B Cell Count | 48,666 | 24 | 11,556 | 1,864 | | 18,347 | 1,082 | 10,343 | 11,625 | 103,507 |
| T Cell Count | 328,608 | 546 | 44,686 | 14,091 | | 151,120 | 2,223 | 4,924 | 142,343 | 688,541 |
| NK Cell Count | 2,915,048 | 15,554 | 1,024,937 | 345,473 | | 1,623,664 | 44,907 | 89,796 | 797,601 | 6,856,980 |
| Neutrophil Count | 11,725,174 | 7,558 | 17,482,860 | 4,461,165 | | 8,139,375 | 408,394 | 1,065,869 | 376,282 | 43,666,677 |
| Total Cluster Count | 2,133,954 | 1,696 | 2,619,122 | 637,081 | | 1,431,314 | 46,829 | 260,719 | 113,163 | 7,243,878 |
| Homotypic CTC Cluster Counts | 50,040 | 4 | 9246 | 365 | | 17,606 | 506 | 69,254 | 1,715 | 148,736 |
| Heterotypic CTC cluster Counts | 33,918 | 8 | 27235 | 3,093 | | 47,239 | 511 | 57,348 | 2,124 | 171,476 |
| Immune Cluster Counts | 2,049,996 | 1,684 | 2,582,641 | 633,623 | | 1,366,469 | 45,812 | 134,117 | 109,324 | 6,923,666 |

### Supplementary Table 2. B06 breast cancer and U08 prostate cancer patient data

| NU16B06 Breast Cancer Patients | Overall |
|---|---|
| **n** | 415 |
| Age at Start (mean (SD)) | 54.75 (12.26) |
| Race (%) | |
| Asian | 16 ( 3.9) |
| Black or African American | 62 (14.9) |
| Unknown | 48 (11.6) |
| White | 289 (69.6) |
| Stage (%) | |
| 3 | 42 (10.5) |
| 4 | 358 (89.5) |
| Subtype (%) | |
| HER2-enriched | 81 (19.5) |
| Luminal A | 157 (37.8) |
| Luminal B | 68 (16.4) |
| Triple-negative | 107 (25.8) |
| Unknown | 2 ( 0.5) |
| CTC Positive (%) | 207 (49.9) |
| Endocrine Therapy (%) | 284 (68.4) |
| Chemotherapy (%) | 375 (90.4) |
| HER2 Targeted Therapy (%) | 154 (37.1) |
| Immunotherapy (%) | 92 (22.2) |
| **NU18U08 Prostate Cancer Patients** | **Overall** |
| **n** | 47 |
| Age (mean (SD)) | 67.15 (10.30) |
| Stage (%) | |
| 2 | 3 (7.1) |
| 3 | 3 (7.1) |
| 4 | 36 (85.7) |
| Gleason Grade Group (%) | |
| 1 | 2 (5.4) |
| 3 | 6 (16.2) |
| 4 | 5 (13.5) |
| 5 | 24 (64.9) |
| CTC Positive (%) | 14 (41.2) |
| Chemotherapy (%) | 17 (37.0) |
| Immunotherapy (%) | 1 (2.2) |
| Endocrine_Therapy (%) | 46 (100.0) |
| Targeted_Therapy (%) | 6 (13.0) |
| Other_Therapy (%) | 1 (2.2) |

### Supplementary Figure S1

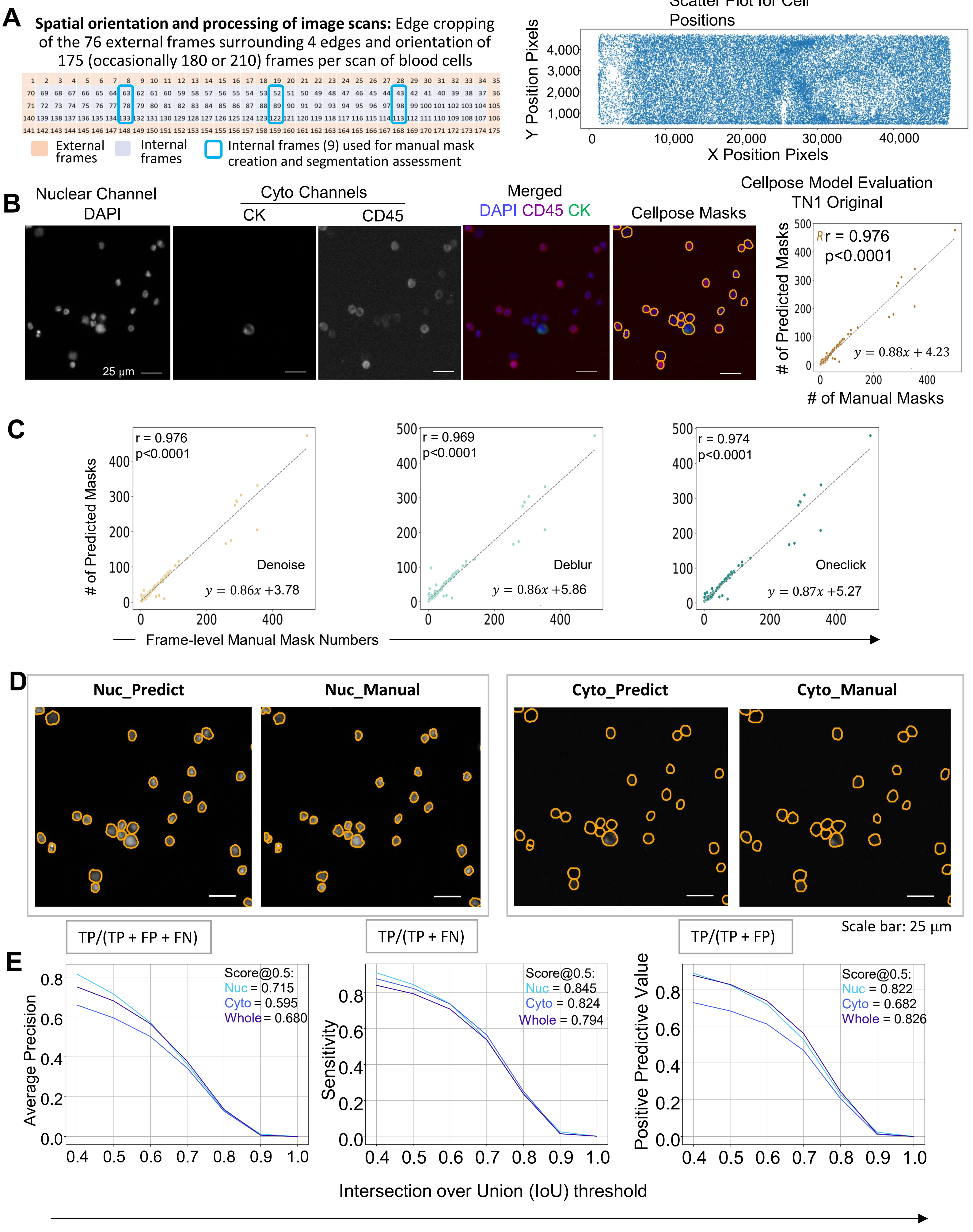

**Supplementary Figure S1. Cell segmentation evaluations**

A. Left penal: The spatial arrangement of 175 frames for the whole scan with annotated frame labels. 9 internal frames were selected for manual mask drawing. Right penal: scatter plot of the center positions for segmented cells throughout the entire scan area from the blood cell image of patient 063-1.

B. Left panels: immunofluorescence images of DAPI (nuclear channel), epithelial cell marker CK and immune cell marker CD45 (Cyto channels), merge channels, and Cellpose-predicted cell masks (whole cell segments). Right panel: Scatter plot and fitted linear equations associating manually labeled cell numbers to Cellpose-predicted cell numbers. The Pearson's correlation coefficient r and the associated p-value are displayed.

C. Scatter plots and fitted linear equations associating manually labelled cell numbers to predicted cell numbers under four pipelines: Original - run Cellpose TN1 model for segmentation directly; Denoise - run Cellpose denoising restoration then TN1; Deblur - Cellpose deblurring then TN1; Oneclick - Cellpose oneclick model (denoising + deblurring) then TN1.

D. Representative image crops for channel-specific raw immunofluorescence images and corresponding cell segmentations. DAPI: DAPI channel for cell nucleus; CK: CK-PE channel for circulating tumor cells; CD45: CD45-APC channel for white blood cells; Nuc_P: model segmentation for nucleus signal (DAPI channel); Nuc_M: manually labelled masks for nuclei; Cyto_P: model segmentation for cytoplasm signal (CK + CD45); Cyto_M: manually labelled masks for cytoplasms; All_P: model segmentation for the whole cell signal.

E. Line plots for average precision, sensitivity and positive predicted value versus multiple thresholds for Intersection over Union (IoU). The segmentation was evaluated from three perspectives - Nucleus, Cytoplasm and the Whole cell. TP-True Positive; FP-False Positive; FN-False Negative.

### Supplementary Figure S2

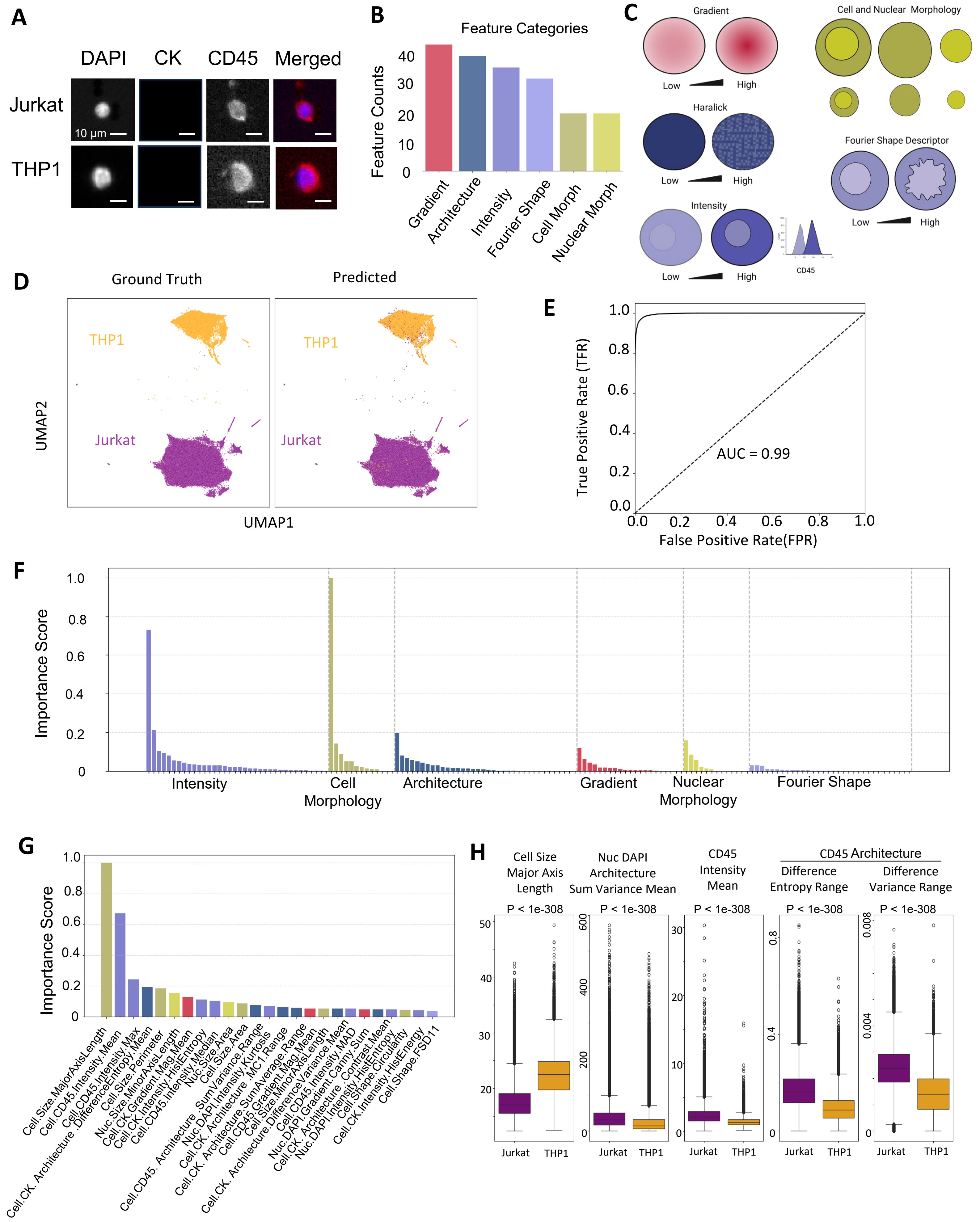

**Supplementary Figure S2. Using machine learning/deep learning to differentiate blood cell types**

A. Image of THP-1 (monocytic) and Jurkat (lymphocytic) cells in DAPI and CD45 channels
B. Counts of cell image features in six categories, including signal gradient (44), Architecture (40), intensity (36), fourier shape (32), cell morphology (20), and nuclear morphology (20), which are used in CCIP for machine learning, Xgboost, to differentiate cells and identify cell types.
C. Schematic of image features extracted within the above six categories, with simulated low and high signals.
D. Supervised UMAPs of 80% of the ground truth immune cells (THP1 and Jurkat) and predicted labels from 20% of the isolated cells, which were loaded to the CellSearch platform for immunofluorescence staining and imaging (CD45, DAPI, CK, and HER2).
E. Receiver Operating Characteristic (ROC) curves with true positive rate (TPR) against false positive rate (FPR) for performance evaluation of the cell image feature machine learning-based CCIP model in predicting THP1 and Jurkat cells, and the diagnostic ability of binary classifiers.
F. Importance score of all features by six categories used in CCIP to identify THP1 and Jurkat cells.
G. Importance score of a list of the top 25 features used in Xgboost-based CCIP to identify two cell types (THP1 and Jurkat cells)
H. Bar graphs of the top 5 features showing the differences between Jurkat and THP1 cells using Student's T-test to find the p-value (n=306,009 Jurkat cells and 110,750 THP1 cells).

### Supplementary Figure S3

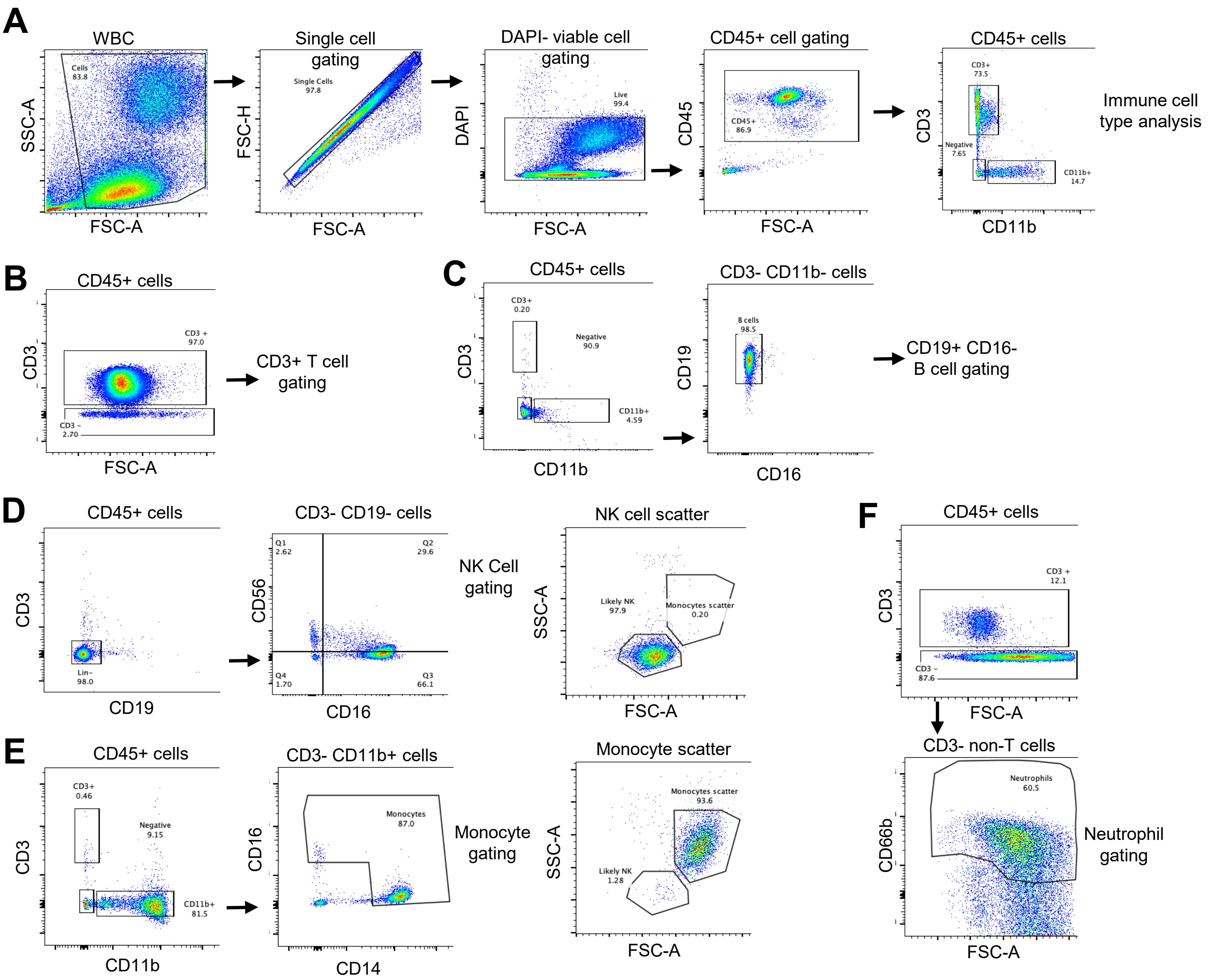

**Supplementary Figure S3. Flow Cytometry panels from bead isolated T Cells, B Cells, NK cells, and Monocytes and ficoll isolated neutrophils used for CCIP model training**

A. Representative flow cytometry dot plot of total white blood cells (WBCs), illustrating the gating strategy for single cells, viable cells (DAPI-), and CD45+ cells, used to identify the subsequent lineage of immune cells, such as T cells (CD3+) and myeloid populations (CD11b+)
B. Flow cytometry panels of bead-isolated T cells (CD45+CD3+, 97.0%) used for CCIP model training
C. Flow cytometry panels of bead isolated B cells (CD45+CD19+CD3-CD11b-, >90%) used for CCIP model training
D. Flow cytometry panels of bead isolated NK cells (CD45+CD3-CD19- CD16+CD56- or CD16-CD56+, >95%) used for CCIP model training
E. Flow cytometry panels of bead isolated Monocytes (CD45+CD3-CD11b+CD14+/CD16+) used for CCIP model training
F. Flow cytometry panels of Ficoll separated neutrophils (CD45+CD3-CD66b) used for CCIP model training

### Supplementary Figure S4

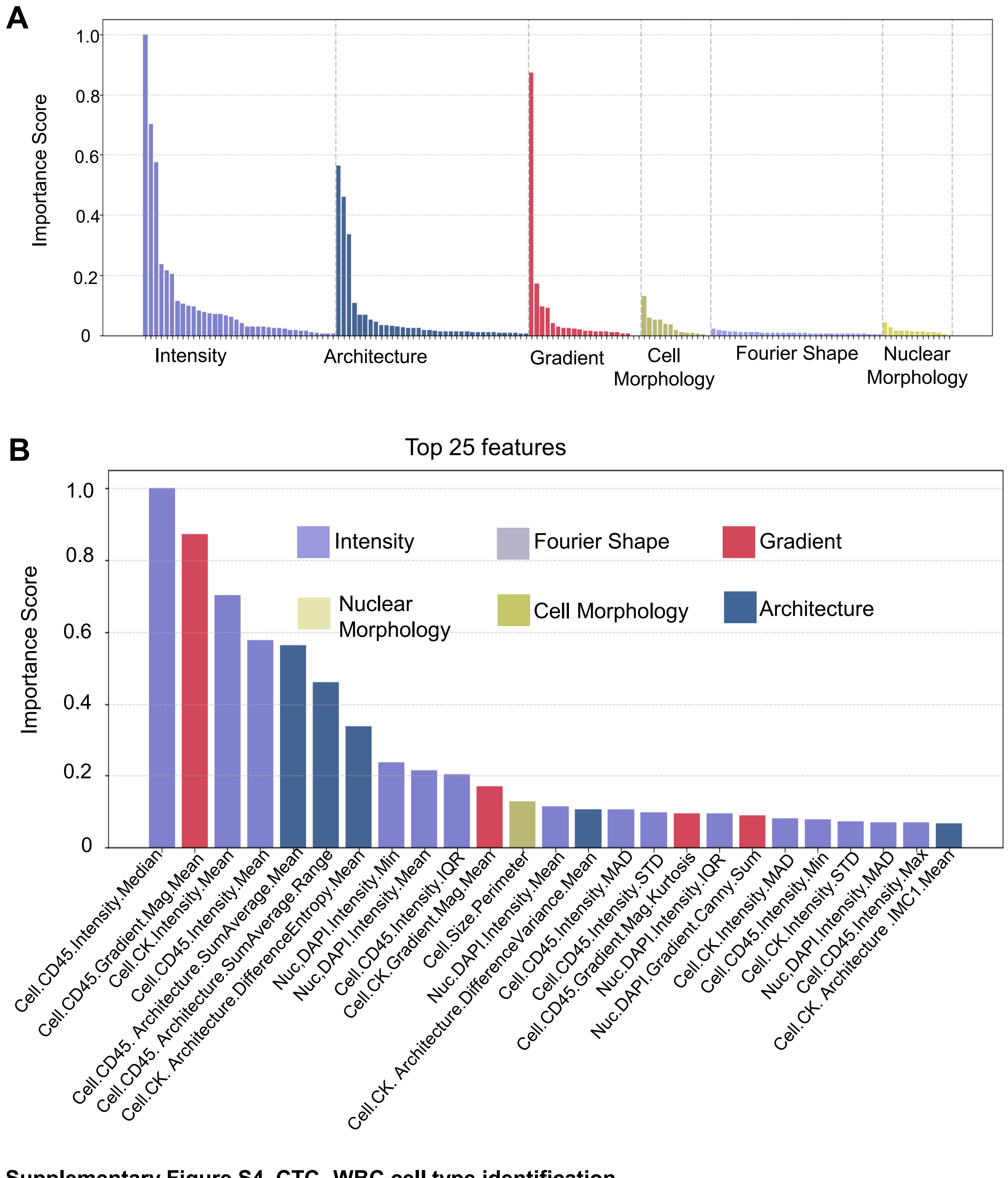

**Supplementary Figure S4. CTC- WBC cell type identification**

A. Feature analysis from the trained classification model depicting how each of the categories are utilized in cell type prediction utilizing normalized importance scores
B. 25 most important features based on normalized importance scores and Feature importance analysis from the trained classification model.

#### Supplementary Figure S5

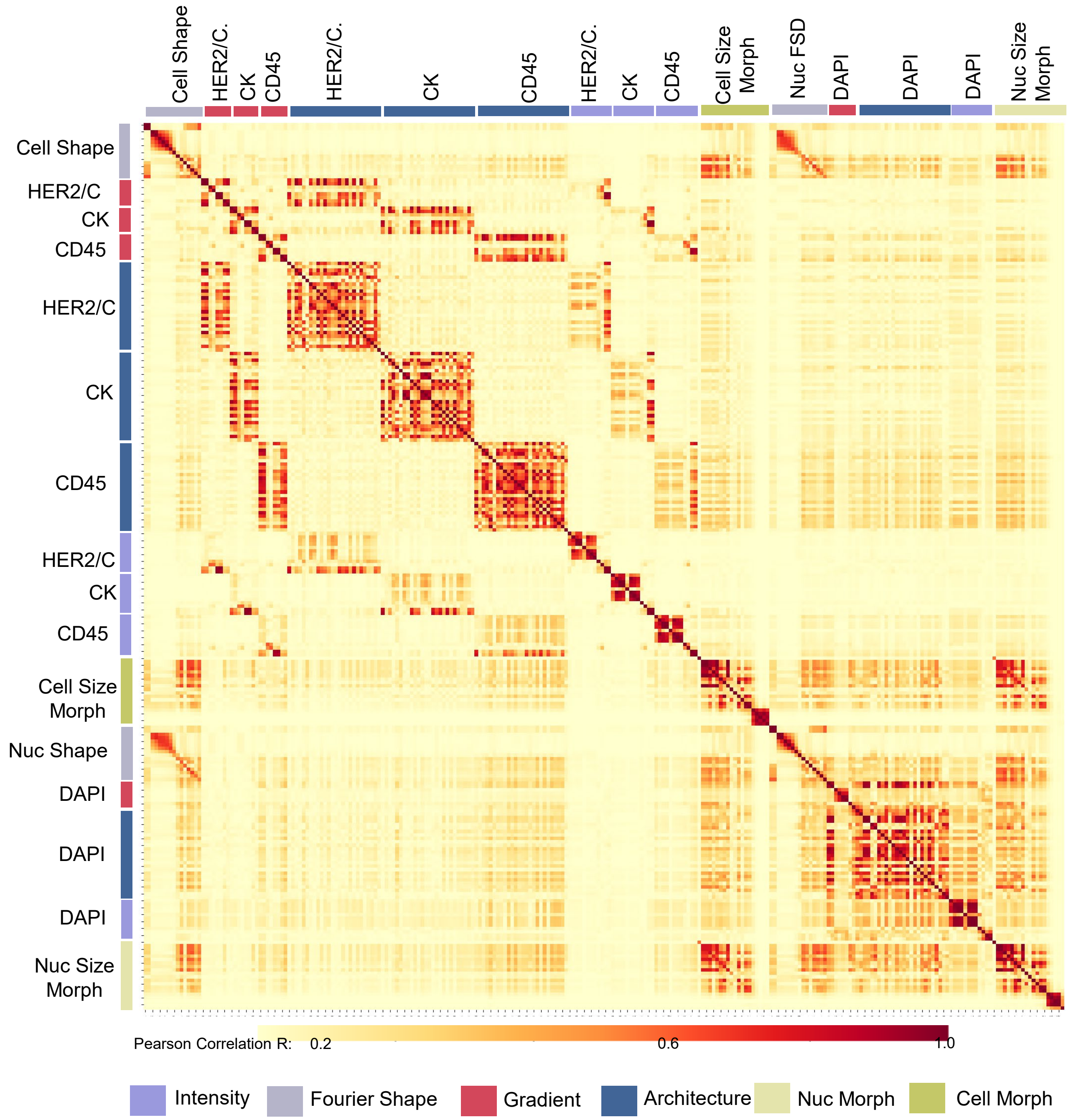

**Supplementary Figure S5. Heatmap of all image features,** depicting Pearson's R correlation of features used for cell type prediction after analyzing 53,871,951 cells from all the scans

#### Supplementary Figure S6

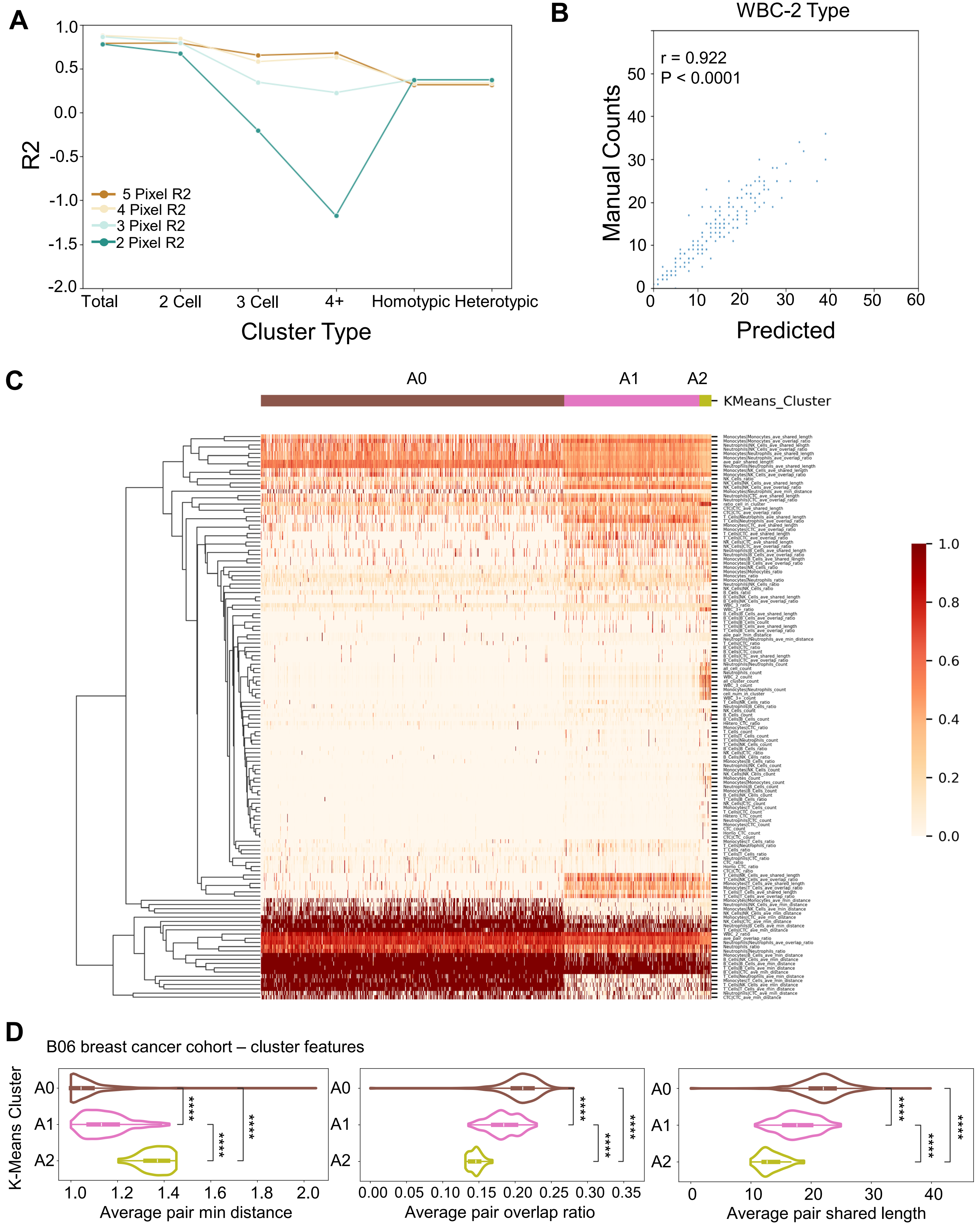

**Supplementary Figure S6. Supporting data for cell cluster detection and unsupervised patient stratifications.**

A. Selection of the optimal mask dilation strength (number of pixels) for pairwise cell contact determination. Four and five-pixel numbers show comparable performance (R-squared value) for all cluster types, but we chose four pixels for dilation as it shows slightly better results for CTC clusters.
B. The scatter plot comparing computationally detected cell cluster numbers with manual counts for white blood cell (WBC) clusters of size two. Pearson correlation coefficient with associated p-value were calculated between two sets of data.
C. Cluster heatmap for the complete list of cell/cluster features over three abundancy groups of B06. Features are normalized into values between 0 and 1. The dendrogram from hierarchical clustering over features indicates distinct patterns across three groups. Scan columns are grouped by the K-means clustering result.
D. The Violin plots showing distinct patterns across three abundancy groups of B06 scans (A0, A1 and A2) for cell interaction scores (from left to right: the number of pixels for minimum distance between boundaries of two contacting cells; The ratio of intersection over union after mask dilation; The length of shared boundary of two contacting cells in pixels). Mann-Whitney U test was conducted across every two groups with star annotation (*: $0.01 < p <= 0.05$; **: $0.001 < p <= 0.01$; ***: $0.0001 < p <= 0.001$; ****:$p <= 0.0001$).

### Extended Figure S7

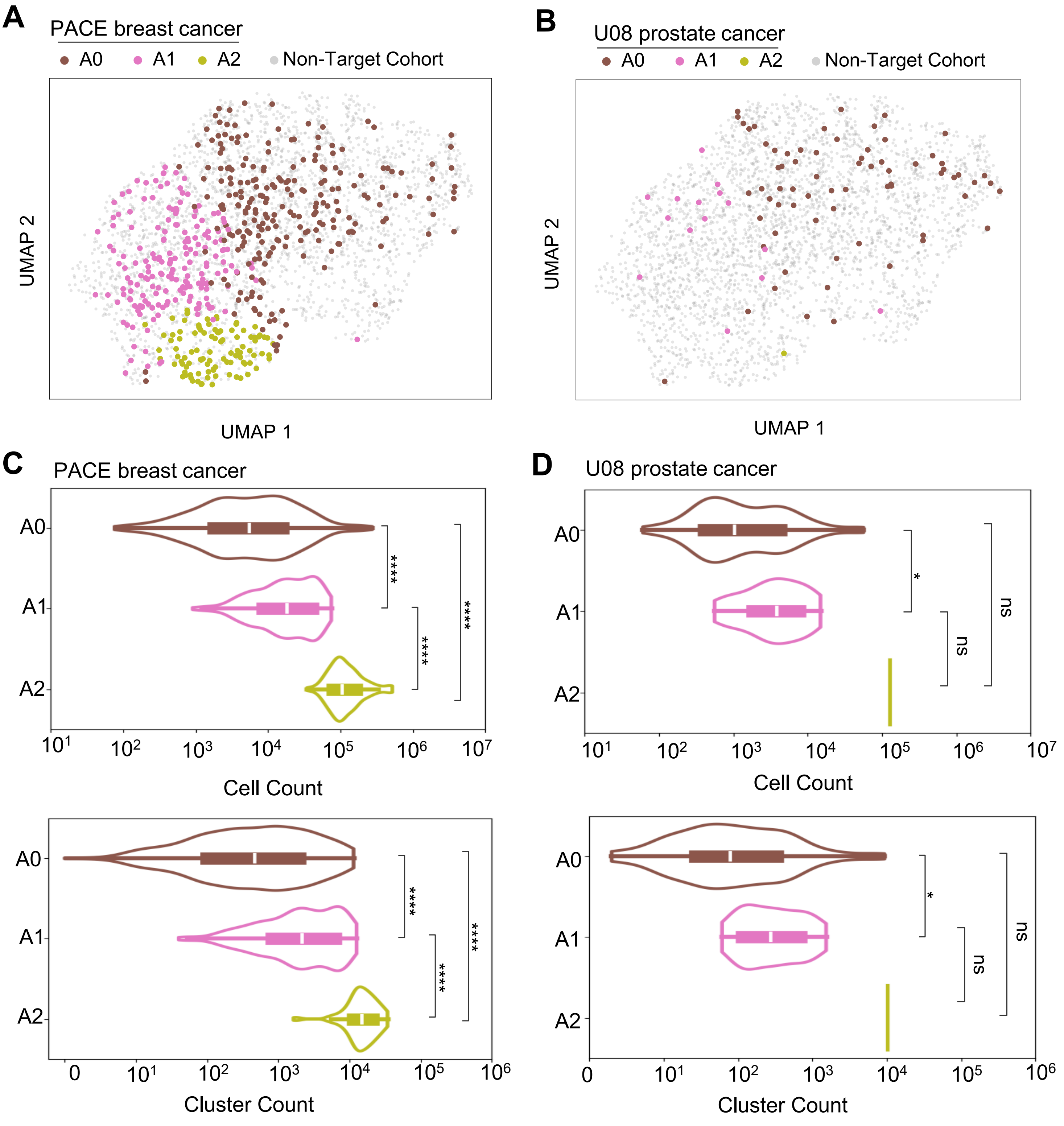

**Extended Figure S7. UMAP of K-means groups for additional breast cancer and prostate cancer cohorts** with respective overall image features.

A. UMAP showing sample distribution for the PACE breast cancer cohort (n=436 scans/tests)

B. UMAP showing sample distribution for the U08 prostate cancer cohort (n=81 scans/tests).

C. Violin plots depicting cell count and cluster count across K-means groups in the PACE breast cancer cohort

D. Violin plots depicting cell count and cluster count across K-means groups in the U08 prostate cancer (NU18U08) cohort.

### Supplementary Figure S8

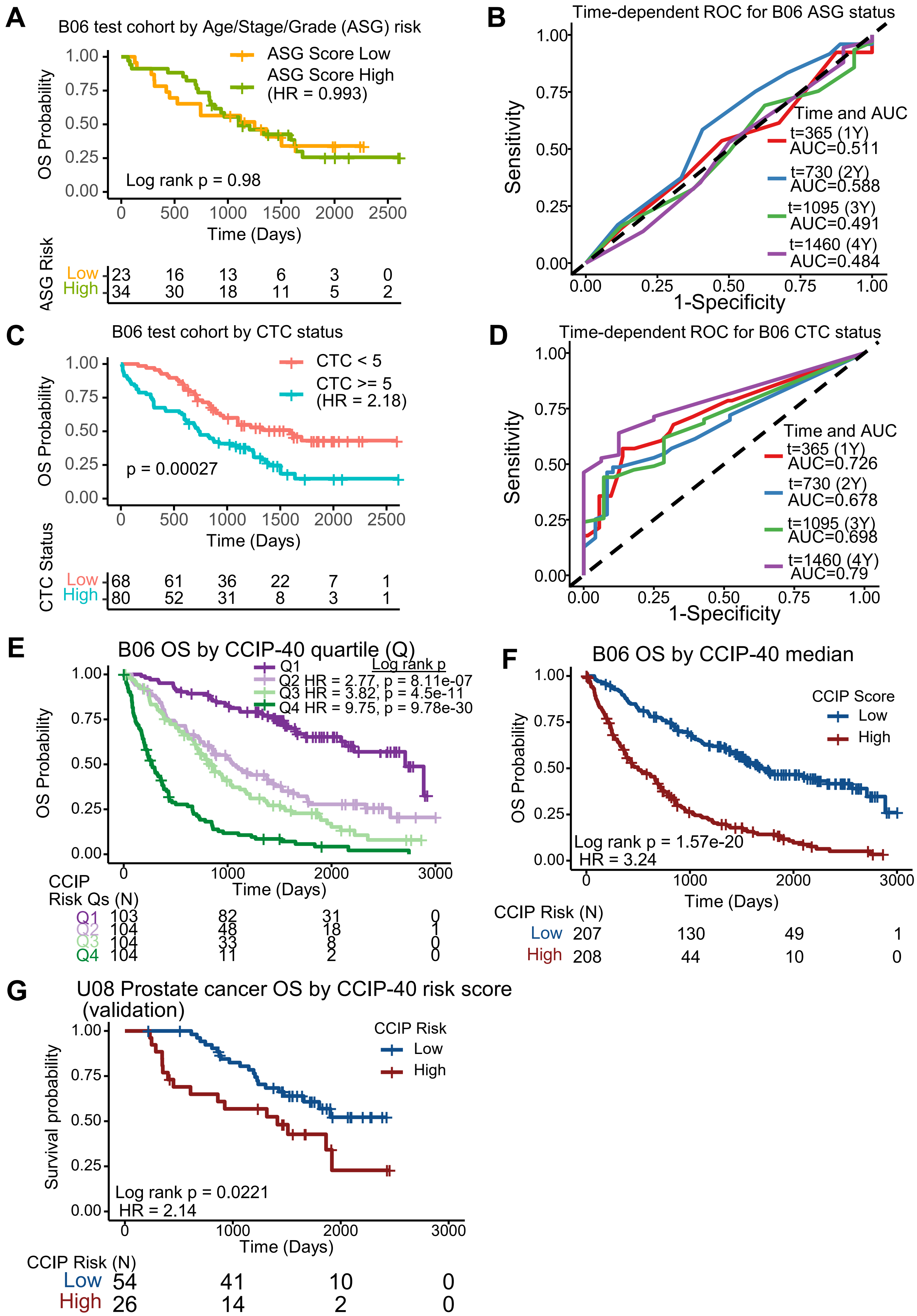

**Supplementary Figure S8. CCIP-40 image risk factors of cells and clusters predict patient outcomes**

A. Kaplan-Meier survival curve using ASG of B06 test data (n = 56 patients). Score weights and stratification were generated using the B06 training data subset. HR=0.99, 95% CI (0.51 to 1.93), log rank p =0.98.
B. Time-Dependent ROC curve for the ASG scores as plotted in A
C. Kaplan-Meier survival curve using CTC positivity as assayed by traditional CellSearch methodology to stratify patients in the B06 test subset. HR=2.18, 95% CI (1.42 to 3.34), log rank p =0.0003.
D. Time-Dependent ROC curve for CTC status as plotted in C
E. Kaplan-Meier curves of overall survival for four quartile groups of B06 breast cancer patients (N=415), stratified based on the CCIP-40 image feature risk score (Extended Fig 8), with the lowest for 1$^{st}$ quartile, Q1, and the highest for 4$^{th}$ quartile, Q4. A Cox proportional hazards model was created for prediction of patient survival. Compared to Q1, Q2 HR=2.77, 95% CI (1.85-4.14), log rank p=8.11e-07; Q3 HR=3.82, 95% CI (2.57-5.69), log rank p=4.5e-11; Q4 HR=9.75, 95% CI (6.58-14.46), log rank p=9.78 e-30.
F. Kaplan-Meier curves of overall survival for the B06 breast cancer patients, two-way stratified by the median CCIP-40 risk scores, combining the Q1 and Q2 as low risk, and Q3 and Q4 in (a) as high risk, respectively. High risk group HR=3.24, 95% CI (2.53-4.16), log rank p=1.57e-20.
G. Cross cancer-type CCIP-40 risk score validation with Kaplan-Meier curves of overall survival for the U08 prostate cancer patients (n = 80 image scans), stratified by the identical CCIP-40 cutoff risk scores of B06 breast cancer cohort as in (c), using the same variables and weights. HR=2.14, 95% CI (1.12-4.11), log rank p=0.022.

### Supplementary Figure S9

| CCIP-40 Risk Factor | MV HR (95% CI) | MV p-value |
|---|---|---|
| CTC_Cell.CK.Architecture.ASM.Range | 9.06e+33 (9.25e+25 to 8.87e+41) | 8.16e-17 |
| CTC_Cell.HER2.Architecture.ASM.Range | 1.49e-15 (6.16e-23 to 3.62e-08) | 8.32e-05 |
| Monocyte_Cell.Shape.FSD14 | 1.06e-14 (1.28e-18 to 8.84e-11) | 2.81e-12 |
| CTC_Cell.CK.Architecture.IMC2.Range | 1.34e+11 (33500000 to 5.38e+14) | 1.42e-09 |
| Monocyte_Cell.CK.Architecture.DifferenceVariance.Mean | 2.34e+10 (4970 to 1.1e+17) | 2.32e-03 |
| NK.Cell_Cell.CK.Architecture.IMC1.Mean | 60200000 (47700 to 7.59e+10) | 8.78e-07 |
| Neutrophil_Cell.CK.Architecture.Correlation.Range | 6.83e-08 (5.97e-17 to 78.1) | 1.21e-01 |
| CTC_Nuc.Shape.FSD14 | 5.41e-06 (2.36e-09 to 0.0124) | 2.12e-03 |
| CTC_Cell.HER2.Architecture.IMC1.Range | 1.09e-05 (2.33e-07 to 0.000512) | 5.86e-09 |
| avg_overlap_ratio_B_Cells_T_Cells | 5840 (59.9 to 569000) | 2.06e-04 |
| Monocyte_Cell.Shape.FSD13 | 0.000837 (3.52e-06 to 0.199) | 1.12e-02 |
| T.Cell_Cell.CD45.Intensity.HistEnergy | 220 (21.5 to 2240) | 5.33e-06 |
| T.Cell_Cell.Shape.FSD14 | 87.9 (0.0235 to 329000) | 2.86e-01 |
| Neutrophil_Cell.CK.Architecture.Correlation.Mean | 0.0122 (0.000142 to 1.04) | 5.19e-02 |
| CTC_Cell.HER2.Architecture.IMC1.Mean | 51.9 (17.7 to 153) | 6.97e-13 |
| avg_overlap_ratio_CTC_Neutrophils | 24 (6.55 to 87.8) | 1.60e-06 |
| CTC_Cell.CK.Architecture.Correlation.Range | 0.163 (2.64e-05 to 1000) | 6.83e-01 |
| NK.Cell_Cell.HER2.Intensity.Min | 5.71 (1.64 to 19.8) | 6.10e-03 |
| T.Cell_Cell.CK.Architecture.ASM.Mean | 4.73 (3.14 to 7.11) | 8.96e-14 |
| Monocyte_Cell.CD45.Gradient.Mag.Skewness | 0.246 (0.151 to 0.399) | 1.49e-08 |
| NK.Cell_Cell.CK.Architecture.IMC2.Mean | 0.267 (0.113 to 0.629) | 2.55e-03 |
| NK.Cell_Cell.CK.Intensity.Skewness | 0.275 (0.154 to 0.492) | 1.38e-05 |
| CTC_Cell.CD45.Architecture.Correlation.Range | 0.359 (0.000128 to 1010) | 8.01e-01 |
| Neutrophil_Cell.CK.Gradient.Mag.Skewness | 0.464 (0.273 to 0.787) | 4.43e-03 |
| B.Cell_Nuc.DAPI.Gradient.Mag.Kurtosis | 2.03 (1.41 to 2.92) | 1.34e-04 |
| Monocyte_Cell.CK.Architecture.ASM.Mean | 1.86 (1.25 to 2.75) | 2.10e-03 |
| Neutrophil_Cell.CK.Architecture.SumEntropy.Mean | 0.576 (0.517 to 0.642) | 9.05e-24 |
| CTC_Cell.CK.Intensity.Skewness | 1.67 (1.32 to 2.1) | 1.44e-05 |
| Neutrophil_Cell.CD45.Gradient.Mag.Kurtosis | 0.602 (0.449 to 0.806) | 6.51e-04 |
| comp_B_Cells_B_Cells | 1.25 (1.15 to 1.36) | 2.11e-07 |
| B_Cells_B_Cells | 1.25 (1.15 to 1.36) | 2.11e-07 |
| CTC_Nuc.DAPI.Architecture.Contrast.Mean | 1.23 (1.14 to 1.33) | 4.87e-08 |
| CTC_Cell.CK.Architecture.Contrast.Mean | 0.814 (0.767 to 0.865) | 2.65e-11 |
| CTC_Cell.HER2.Architecture.Contrast.Range | 0.827 (0.761 to 0.898) | 7.18e-06 |
| Monocyte_Cell.CK.Architecture.SumEntropy.Mean | 0.856 (0.774 to 0.948) | 2.68e-03 |
| CTC_Nuc.DAPI.Gradient.Mag.Std | 1.16 (1.1 to 1.23) | 1.53e-07 |
| CTC_Cell.HER2.Architecture.SumVariance.Range | 0.872 (0.814 to 0.934) | 9.13e-05 |
| Monocyte_Cell.CK.Architecture.Entropy.Mean | 0.908 (0.852 to 0.967) | 2.89e-03 |
| CTC_Cell.CK.Architecture.SumAverage.Mean | 0.927 (0.909 to 0.945) | 5.00e-14 |
| Neutrophil_Cell.CK.Architecture.SumVariance.Mean | 0.964 (0.937 to 0.99) | 8.16e-03 |
| Neutrophil_Cell.CK.Gradient.Canny.Sum | 0.993 (0.989 to 0.998) | 5.70e-03 |
| Monocyte_Cell.CK.Gradient.Canny.Sum | 0.995 (0.991 to 0.999) | 7.85e-03 |
| Num_Nphils | 1 (1 to 1) | 9.79e-04 |

**N = 332 patients**

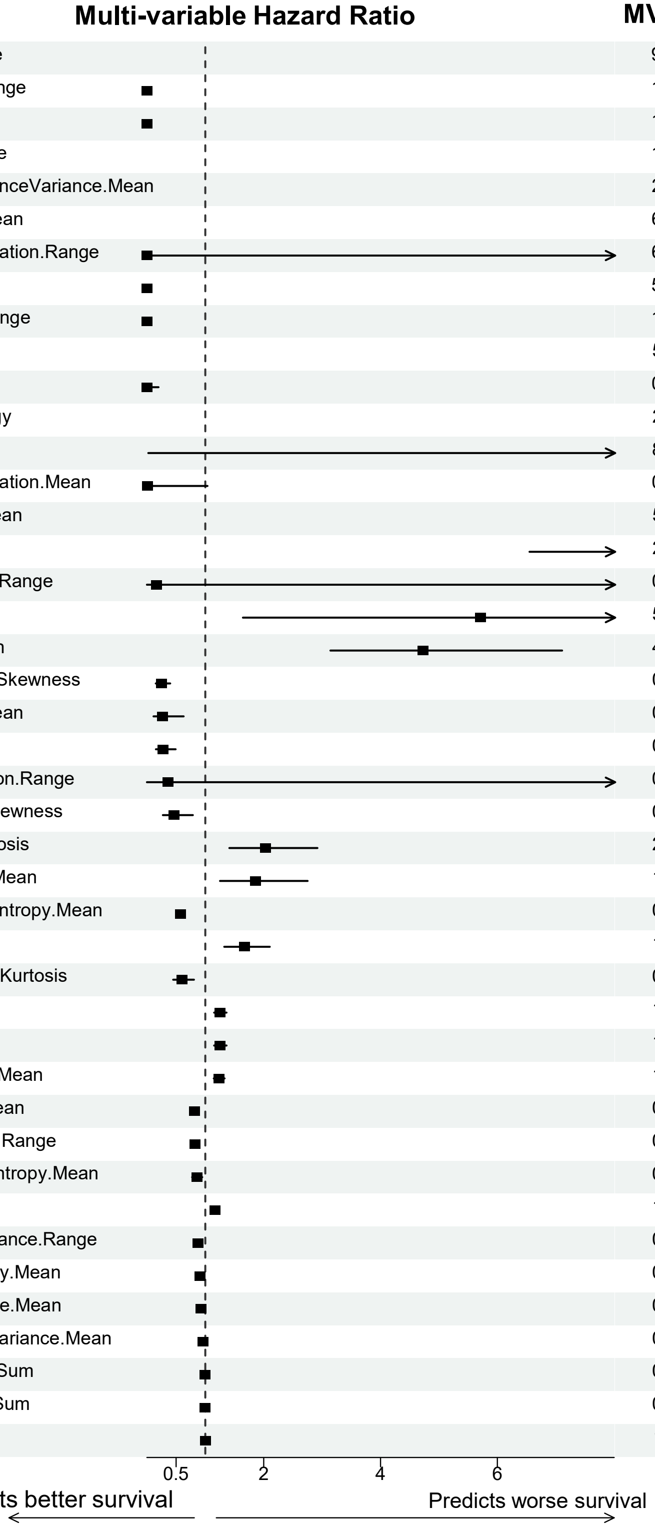

**Supplementary Figure S9. CCIP top 40 image features (cells and clusters) associated with patient overall survival (OS).**

CCIP top 40+ risk factors of cell and cluster image features with proportional hazards for OS, evaluated using LASSO regularization. These features were analyzed using a multi-variable Cox proportional hazards model in which patient demographic features (race and tumor subtype) are not among the top list. The B06 patients (N=415) were divided into 10 groups, with nine used to create a model and the tenth used to generate a risk score. P values adjusted using FDR.

### Supplementary Figure S10

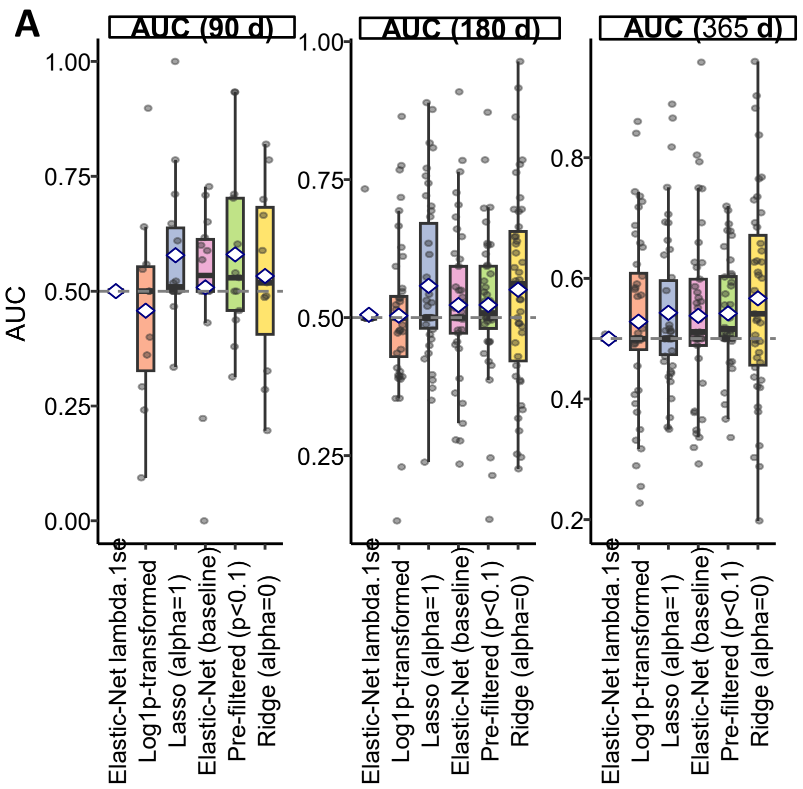

B

| Model | AUC (90 d) | AUC (180 d) | AUC (365 d) |
|---|---|---|---|
| Elastic-Net lambda.1se | 0.500 ± 0.000 | 0.505 ± 0.035 | 0.500 ± 0.001 |
| Log1p-transformed | 0.458 ± 0.216 | 0.504 ± 0.134 | 0.528 ± 0.145 |
| Lasso (alpha=1) | 0.578 ± 0.165 | 0.558 ± 0.148 | 0.543 ± 0.128 |
| Elastic-Net (baseline) | 0.508 ± 0.193 | 0.523 ± 0.140 | 0.538 ± 0.142 |
| Pre-filtered (p<0.1) | 0.580 ± 0.193 | 0.523 ± 0.137 | 0.542 ± 0.092 |
| Ridge (alpha=0) | 0.533 ± 0.204 | 0.551 ± 0.179 | 0.567 ± 0.170 |

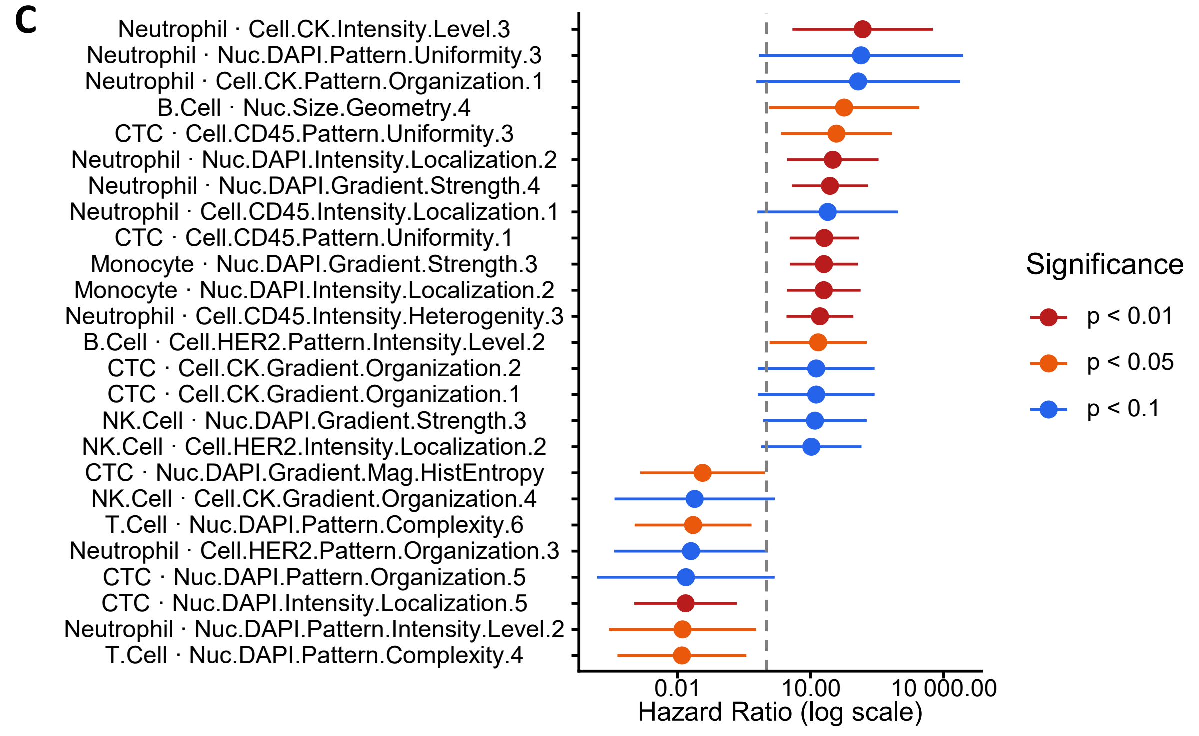

**Supplementary Figure S10. CTC imaging feature model benchmarking and univariate associations with chemotherapy progression-free survival (PFS).**

A. Time-dependent AUC at 90, 180, and 365 days for six penalized Cox regression strategies evaluated by 5-fold cross-validation with 10 repeats. Each point represents one fold; diamonds indicate the cross-validated mean. Dashed line at 0.5 denotes chance performance. Models tested include Elastic-Net (lambda.1se), Log1p-transformed Lasso, Lasso (alpha=1), Elastic-Net (baseline), pre-filtered Lasso (univariate p<0.1), and Ridge (alpha=0). Clinical covariates (molecular subtype, Stage IV) were force-included in all models.

B. Summary table of mean ± SD time-dependent AUC across cross-validation folds for each model.

C. Univariate Cox proportional hazards forest plot showing the top 25 CTC imaging features most strongly associated with chemotherapy PFS (ranked by |log HR|, filtered at p<0.1). Points indicate hazard ratios; horizontal bars indicate 95% confidence intervals. Dashed vertical line at HR=1. Feature names are formatted as Cell Type · Feature Group Number. Color indicates the significance level of the univariate Wald test.

### Supplementary Figure S11

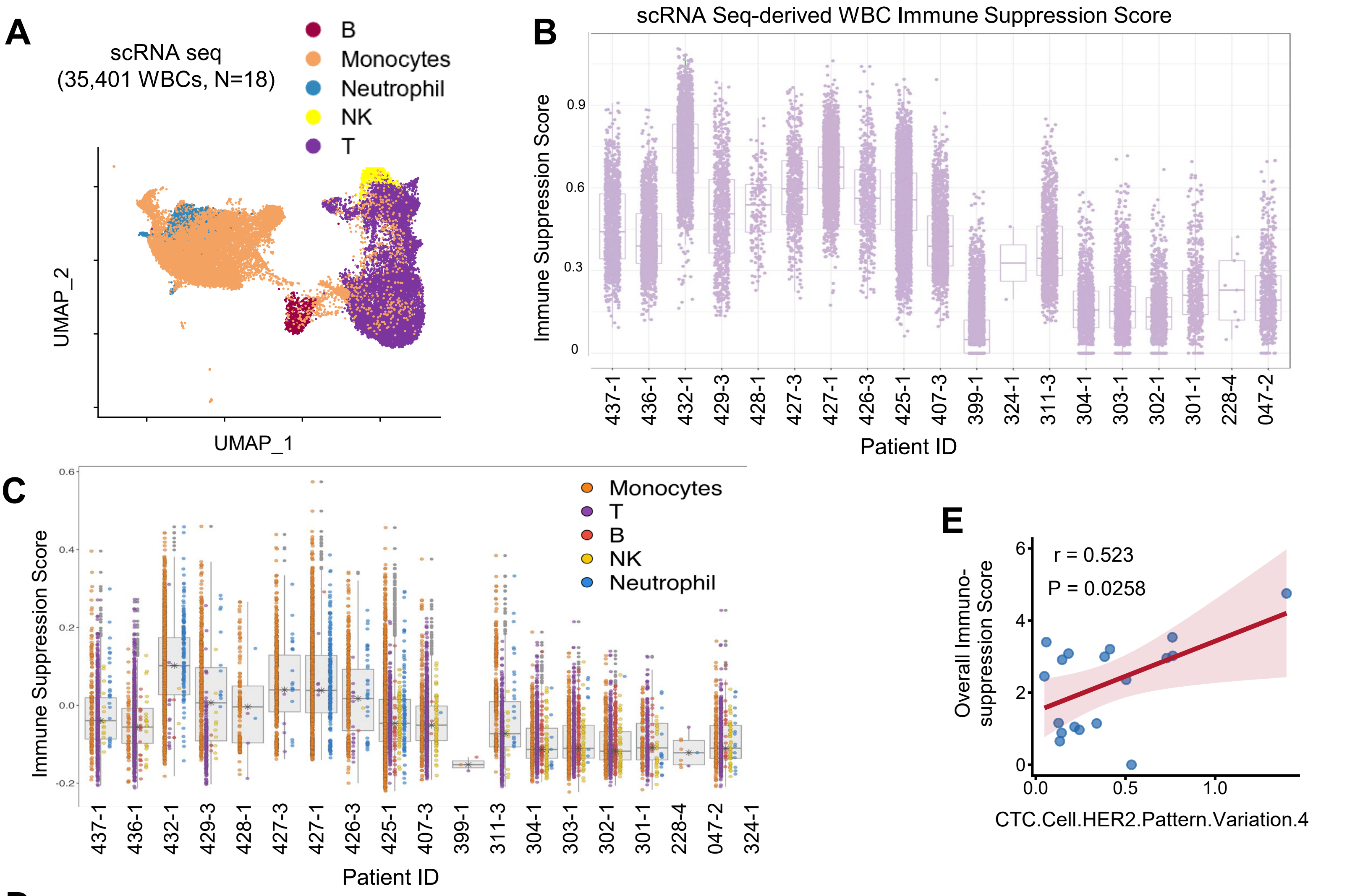

D

| Cell Type of Immune Suppression | Image Feature | HR | HR (95% CI) | P-value | Adj P |
|---|---|---|---|---|---|
| Monocytes | CTC.Cell.HER2.Pattern.Variation.4 | | 5.47 (1.11–27.04) | 0.0385 | 0.0646 |
| Monocytes | B.Cell.Cell.CD45.Intensity.Localization.2 | | 3.84 (1.07–13.77) | 0.0404 | 0.0646 |
| Neutrophil | B.Cell.Cell.CD45.Intensity.Localization.2 | | 8.62 (1.81–41.07) | 0.0099 | 0.0646 |
| Neutrophil | CTC.Cell.HER2.Pattern.Variation.4 | | 8.06 (0.93–69.69) | 0.0571 | 0.0646 |
| NK | CTC.Nuc.Size.Geometry.4 | | 5281.23 (0.74–37808560) | 0.0574 | 0.0646 |
| T | CTC.Nuc.Size.Geometry.4 | | 9480.19 (8.03–11197925) | 0.0144 | 0.0646 |
| Overall | CTC.Cell.HER2.Pattern.Variation.4 | | 7.01 (1.31–37.63) | 0.0258 | 0.0646 |
| Overall | B.Cell.Cell.CD45.Intensity.Localization.2 | | 3.87 (0.95–15.68) | 0.0574 | 0.0646 |
| Overall | NK.Cell.Cell.HER2.Intensity.Level.1 | | 0.01 (0.00–1.51) | 0.0714 | 0.0714 |

0.01 1 100

Associated with Immune Activation Associated with Immune Suppression

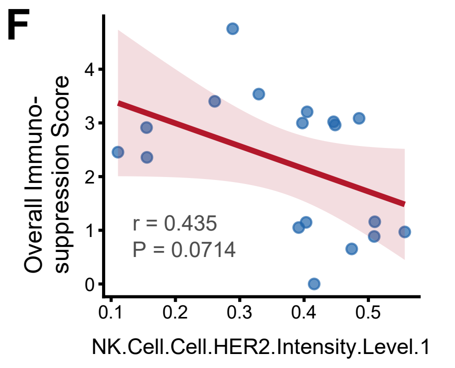

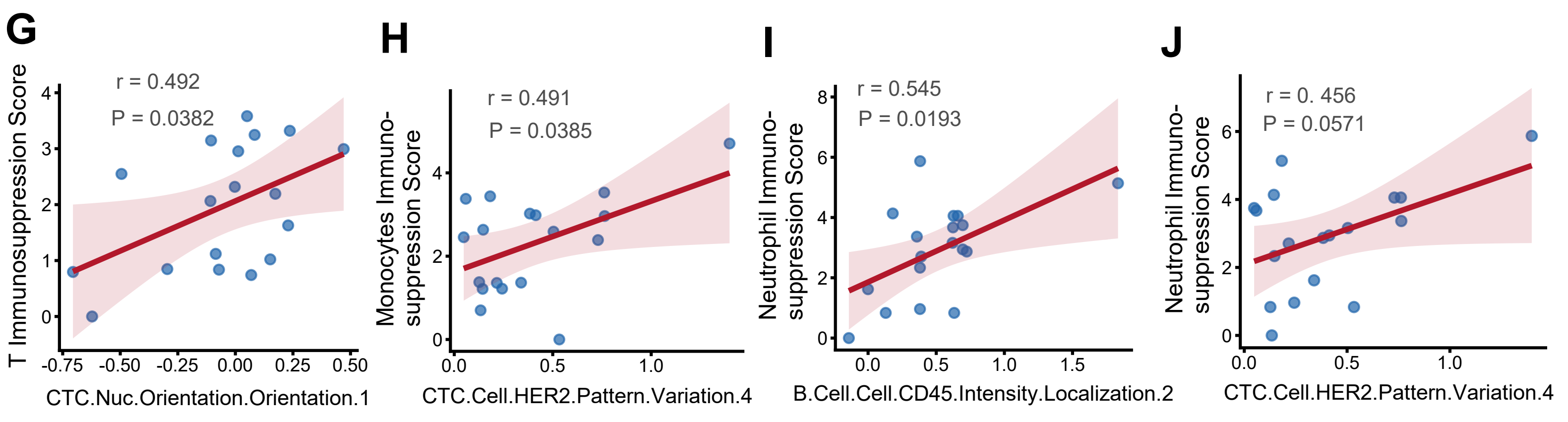

**Supplementary Figure S11. scRNA seq-derived immune suppression scores associated with patient survival-related CCIP image features.**

**A.** UMAP visualization of single-cell RNA sequencing profiles of 35,401 WBCs collected from B06 breast cancer patients (N = 18). Distinct groups represent major immune cell populations, corresponding to the CCIP identification groups of B cells, T cells, NK cells, monocytes, and neutrophils.

**B.** Box plots of the immune suppression score distribution of individual WBCs per patient, with overlaid dots representing individual cells. Immunosuppressive scores were calculated using a predefined gene panel.

**C. Immunosuppressive scores of WBCs with 5 immune cell types**, including monocytes, T, B, NK, and neutrophils, calculated using a predefined gene panel. Box plots show the distribution of scores per patient, with overlaid dots representing individual cells colored by cell type.

**D.** Forest plots showing univariate Cox proportional hazards analysis of all previously identified imaging-derived features (CCIP-14 and therapy response-specific features) associated with immune suppression score. Points represent hazard ratios (HR), and horizontal lines indicate 95% confidence intervals (CI). Features with HR > 1 are associated with higher score, while HR < 1 indicates vice versa. Statistical significance (P- values) is computed by the Wald test and corresponding significance levels.

**E-J.** Scatter plot comparing immune suppression score (overall or one specific immune cell) with image features associated with either OS or post-chemotherapy PFS. The Pearson's correlation coefficient r and associated P value for linear correlation between two are displayed.
